# Epistatic fitness landscapes emerge from parallel adaptive walks in breeding network metapopulations

**DOI:** 10.64898/2026.03.18.712732

**Authors:** Theodore Monyak, Geoffrey P. Morris

**Author notes:** Correspondence should be addressed to Geoffrey P. Morris.

## Abstract

Global networks of crop breeding programs leverage diverse germplasm, but diversity increases the complexity of maintaining stability in their elite genepools. To characterize genetic heterogeneity in breeding metapopulations and develop insights on how to manage it, we simulated the evolution of breeding populations on fitness landscapes. We revealed the geometric decrease in the average effect size of alleles segregating as standing variation that become fixed along an adaptive walk. We also demonstrated how independent adaptive walks of subpopulations are influenced by genetic drift, leading to cryptic genetic heterogeneity among elite genepools. This variation is released when elite lines derived from independent subpopulations are crossed, leading to segregation for 2-4X more major QTL in admixed families as in unadmixed families, and 2-4X more epistatic interactions. The emergent property of fitness epistasis for traits under stabilizing selection is well-understood in evolutionary genetics, but under-appreciated in crop quantitative genetics. To highlight the importance of this phenomenon, we constructed an empirical genotype-to-fitness landscape from the sorghum NAM, a global admixed prebreeding resource, demonstrating the utility of fitness landscapes for inferring genetic compatibilities within metapopulations. Our findings suggest that in breeding networks, strategies for effective germplasm exchange must account for epistasis in the oligogenic component of the genetic architecture of locally-adapted traits.

**Article summary:** Modern public sector crop improvement happens in networks of breeding programs that routinely exchange genetic information. Traditional models for understanding quantitative traits have limited predictiveness in situations with such genetic heterogeneity. This study uses breeding simulations and empirical data to show the utility of the fitness landscape framework for characterizing the genetic architecture of complex traits in breeding metapopulations. By simulating the evolution of breeding programs and integration into networks, it demonstrates how epistatic interactions between large-effect alleles are a fundamental property that must be accounted for when exchanging germplasm.

**Graphical Abstract:** 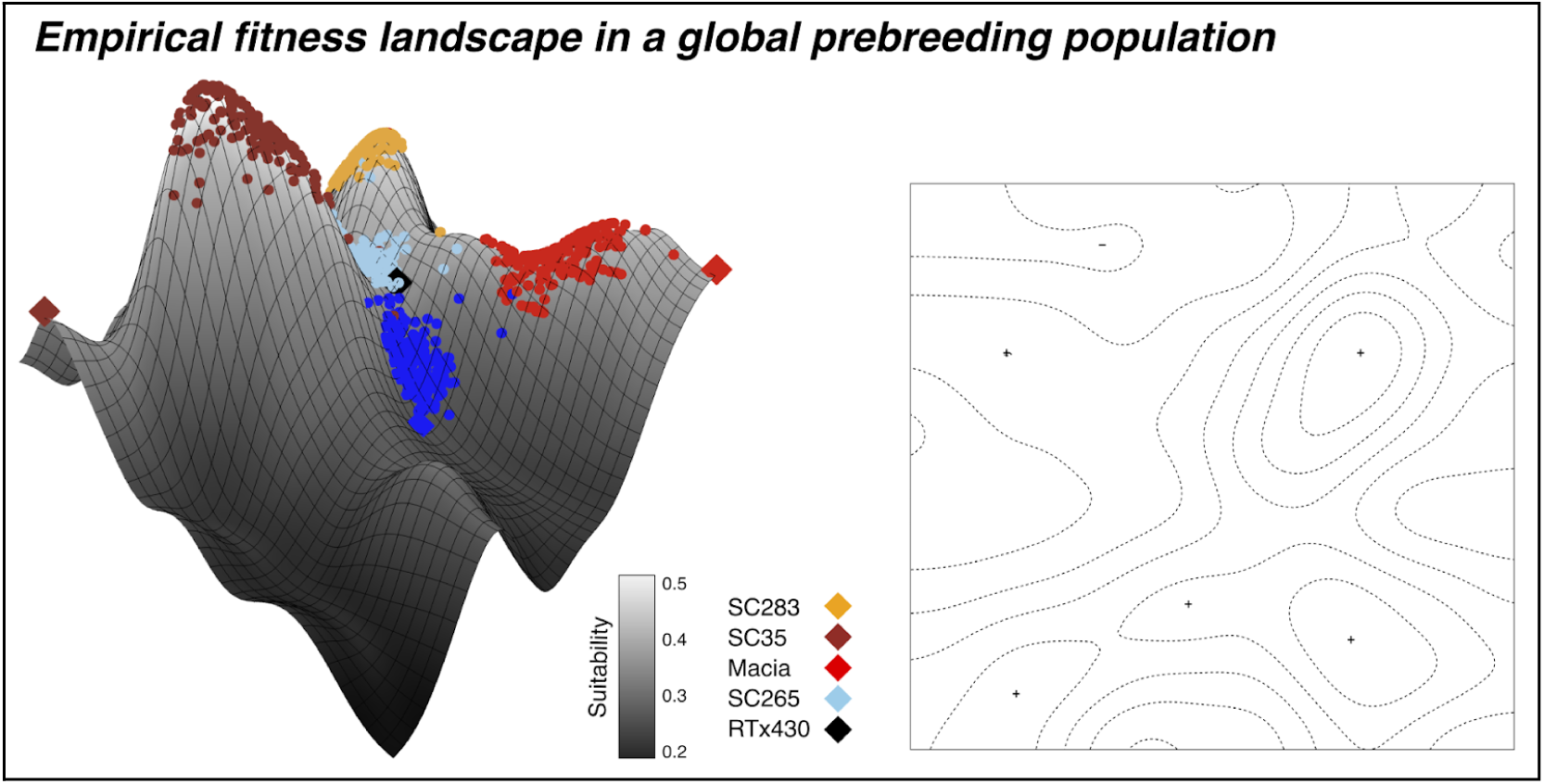

## INTRODUCTION

Genetic theory continually advances based on observations from plant and animal breeding (Darwin 1876; Fisher 1930; Wright 1931; Lande and Thompson 1990), which in turn leads to improved breeding strategies reflecting a more complete understanding of genetics. For example, population genetics knowledge informs our understanding that genetic gain in breeding depends on a balance between maintaining elite genetic homogeneity and exploiting genetic diversity (Allard 1999; Hallauer et al. 2010; Bernardo 2014; Swarup et al. 2021). Historically, this has been achieved with closed breeding populations and the periodic introgression of beneficial alleles (Holley and Goodman 1988; Tanksley and McCouch 1997; Cobb, Juma, et al. 2019). In contrast, modern networks of public sector breeding programs routinely leverage information across diverse germplasm to broaden the genetic diversity of their breeding populations, rapidly respond to novel agronomic stresses, and advance regional breeding objectives (Atlin et al. 2017; Muleta et al. 2022; Maina et al. 2025; Morris et al. 2026). This approach is particularly being adopted by nascent public sector breeding programs in sub-Saharan Africa managed by the national agricultural research systems (NARS) and the Consultative Group on International Agricultural Research (CGIAR) (Kholová et al. 2021; Das et al. 2024). Success of breeding networks will require a reconsideration of the standard genetic assumptions underlying crop improvement methods.

Greater exchange of germplasm within a breeding network heightens the challenge of maintaining stability in elite genepools. Breeders typically make elite-by-elite crosses (Fehr 1984; Allard 1999; Rutkoski 2019; Bernardo 2020), which is expected to produce progeny with parental phenotypes when the parents are isogenic at the major loci underlying those traits (i.e. iso*-*elite) (Fall et al. 2026). However, if the parents have contrasting haplotypes at those loci (i.e. are allo-elite), their progeny may exhibit excessive transgressive segregation, in which their phenotypic range extends well above and below the parental trait values (Rieseberg et al. 1999; Fall et al. 2026). While transgressive segregation is crucial for genetic gain of traits under directional selection (Mackay et al. 2021) such as yield (i.e. *desired* traits), it is undesirable for traits under stabilizing selection (i.e. *attained* traits) (Fall et al. 2026). Transgression can occur when the formation of novel allelic combinations releases cryptic genetic heterogeneity (Hermisson and Wagner 2004; Paaby and Rockman 2014), leading to the dispersal of complementary additive quantitative trait loci (QTL) (Vega and Frey 1980; deVicente and Tanksley 1993; Rieseberg et al. 2003; Castro et al. 2008), rare recessive alleles becoming homozygous (Rick and Smith 1953), or unfavorable epistatic interactions between loci (Li et al. 2015; Soyk et al. 2017; Olatoye et al. 2020).

The genetic architecture of complex traits in a breeding network may be elucidated with shifting balance theory (SBT) and fitness landscapes (Wright 1932; Kauffman and Levin 1987; Whitlock et al. 1995; Fragata et al. 2019). A fitness landscape in the context of crop improvement is a rugged surface of fitness peaks and valleys shaped by agroclimatic and cultural constraints (Allard 1999; Podlich and Cooper 1999; Hammer et al. 2006; Stewart 2008; Messina et al. 2011; Technow et al. 2021) (Supplementary Table 1). A breeding network in this framing constitutes a metapopulation, a group of partially isolated sub-populations, which traverses the landscape through selection and genetic drift (Wright 1931; van Heerwaarden et al. 2010; Cooper et al. 2014; Technow et al. 2021; Technow et al. 2026). Historically, a lack of empirical evidence of population subdivision contributed to skepticism of SBT (Coyne et al. 1997; Crow 2008) but modern genomic studies have revealed that hierarchical population structure is a near-ubiquitous characteristic of crops (Jordan et al. 2005; Jin et al. 2010; Blair et al. 2012; Morris et al. 2013; Xiong et al. 2016), even among obligate outcrossers such as maize (Hufford et al. 2012) and pearl millet (Serba et al. 2019). Further, parallel evolution is expected to produce genetic heterogeneity and interactions among large-effect QTL (Wright 1931; Wood et al. 2005; Blount et al. 2018; Yeaman 2022), especially when there is a low effective population size (Wright 1931), which is characteristic of inbred agricultural crops.

In this study, we sought to reconcile the alternative viewpoints of adaptation in crop breeding by establishing a connection between historical selection of subpopulations and the genetic architecture of complex traits in admixed breeding germplasm. We used *in silico* breeding simulations to construct a trait-to-fitness landscape and investigate the dynamics of adaptive walks of crop populations. Our simulations were representative of the history of an archetypal nascent breeding program: progressing from a small, heterogenous population of landraces to a homogenous elite germplasm through purification of attained traits and selection for higher yield. Applying the same selection pressure to independent populations resulted in stochastic adaptive walks and phenotypic parallelism. We demonstrated that fitness epistasis is an emergent property of this process, underlying the cryptic genetic heterogeneity that is released through the integration of different breeding populations. The approach developed here provides a practical theoretical and empirical framework for use of fitness landscapes to manage breeding metapopulations and develop cross-population breeding strategies.

## MATERIAL AND METHODS

### Simulation of a founder population

For each simulation run, an initial founder population of 2000 biallelic, diploid organisms was simulated in the R package “AlphaSimR” (Gaynor et al. 2021) with coalescent MaCS software (Chen et al. 2009). Each individual had 10 chromosomes each with a genetic length of 100 cM and 1000 segregating sites. 3 traits were simulated: Traits 1 and 2 represented predominantly oligogenic *attained* traits (under stabilizing selection; e.g. flowering time and plant height), and Trait 3 represented a polygenic *desired* trait (under directional selection; e.g. yield potential) (Fall et al. 2026). The initial mean population genetic values of the attained traits were 2 units from the optimum, genetic variance were 0.4 units squared, narrow-sense heritability (*h*^2^) was 0.1, and the number of effective QTL per attained trait, *L,* was 10, 20 or 50, depending on the simulation parameters. These values were selected so that the founder population began simulation at the base of the fitness peak. The initial mean genetic value of Trait 3 was 50 units, with a genetic variance of 50 units squared, (*h*^2^) narrow-sense heritability of 0.05, and 500 QTL throughout the genome. These parameters were chosen to generate a highly polygenic trait with high phenotypic variance and low heritability, consistent with standard assumptions about the genetic basis of yield potential (Meuwissen et al. 2001; Heffner et al. 2009; Bernardo 2020). Each of the traits had an additive genotype-to-trait mapping. Trait values were calculated as the summation across all QTL of its effect size, multiplied by the allelic dosage, plus normally distributed error, as determined by each trait’s specified heritability.

### Defining a geometric series of additive substitution effect sizes

Allele substitution effect sizes for the attained traits in the founder population were defined by the geometric distribution

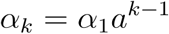

where the substitution effect of the *k*th locus *α_k_* was determined based on *α_1_*, the effect size of the largest-effect QTL, and *a*, the relative size of each subsequent QTL (Lande and Thompson 1990). The parameter *a* was calculated as a function of *L* (Bernardo and Yu 2007), as

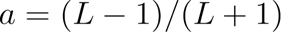

Genome locations for each QTL were randomly selected from the segregating sites in the genome with a minor allele frequency above 0.3, with an equal number of QTL per chromosome. The scaling term *α*_1_ was calculated with the rescaleTraits() function from AlphaSimR for each founder population to give them a consistent initial additive variance. Effect sizes were randomly selected to be positive or negative and recentered to have consistent initial mean trait values. Three different founder trait genetic architecture scenarios were used throughout the simulations: a “low-end” oligogenic scenario (*L* = 10, *a* = 0.82), an “intermediate” oligogenic scenario (*L* = 20, *a* = 0.90), and a “high-end” oligogenic scenario” (*L* = 50, *a* = 0.96).

### Defining a trait-to-fitness landscape

A fitness landscape was defined with a single adaptive ‘suitability’ (*S*) function based on the values of the two attained traits. Suitability measured the adaptedness of an individual in the range of [0,1] to a set of selection constraints, based on the difference between each of the attained trait phenotypes and the optimal phenotype of 0. This fitness peak was modeled with the following Gaussian function, in accordance with literature (Lande 1980; Kimura 1983; Orr 1998; Martin et al. 2007) (Supplementary Table 2), where t_1_ was the value of Trait 1, and t_2_ was the value of Trait 2:

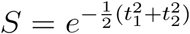

During the ‘breeding’ phase of the simulation, individuals were selected according to their suitability. During the ‘landrace’ phase of the simulation, individuals were selected according to breeding fitness (*w*), meant to represent realized yield, calculated as polygenic desired trait t_3_, penalized by lower suitability scores:

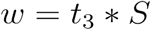

Our trait-to-fitness landscape was rendered based on the suitability function (Fig. 1a), using the R package “plotly” (Sievert 2020).

**Figure 1.**
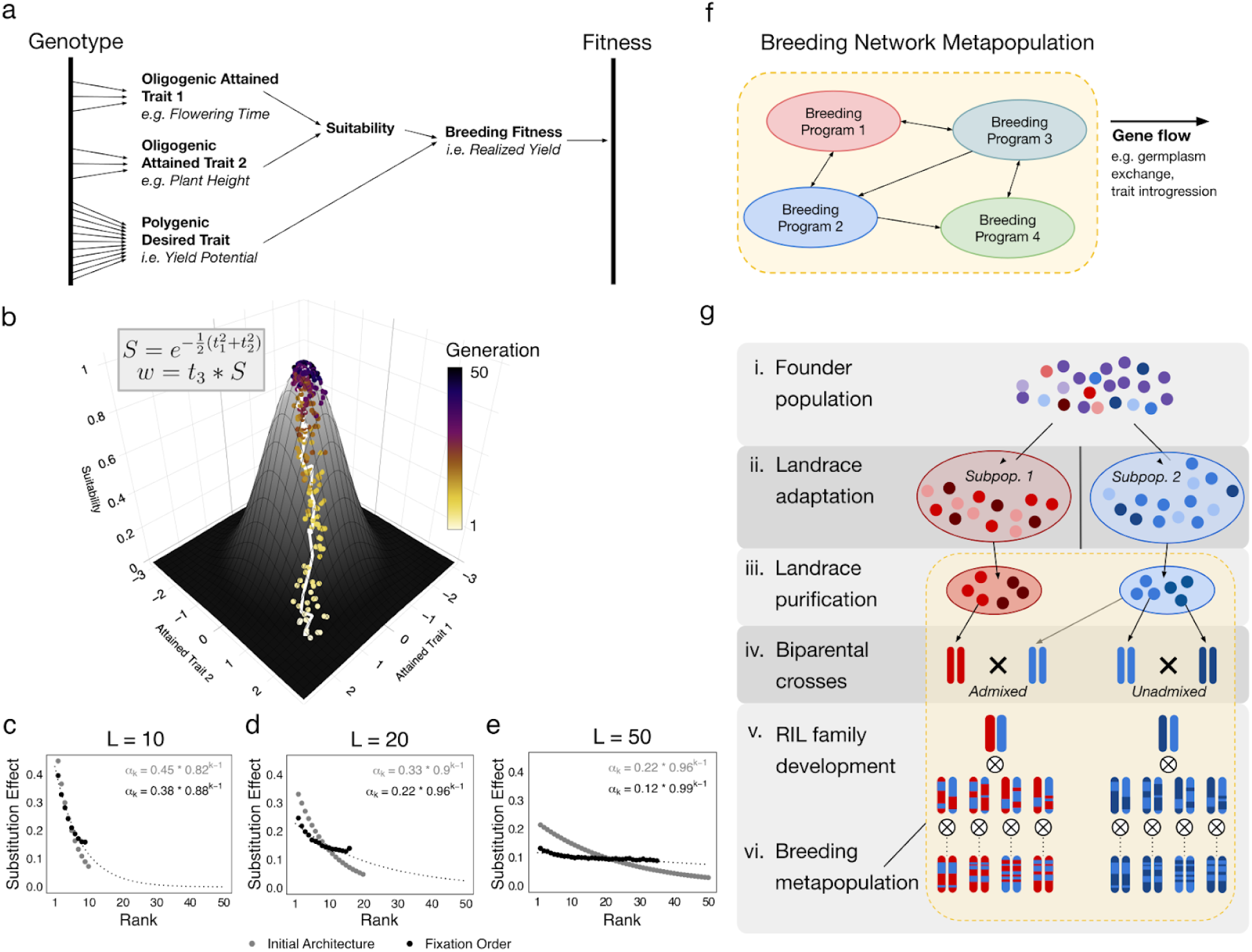
Adaptive walks on a trait-to-fitness landscape lead to the fixation of alleles of progressively smaller effect sizes. (a) A schematic of the genotype-to-fitness map used in our simulations. (b) Our [suitability] trait-to-fitness function produced a single-peaked 3D surface. A population of 1000 individuals was selected for 50 generations according to *w* (breeding fitness), where at each generation, the top 10% of individuals were randomly crossed. 5 individuals from each generation, which were closest to the population means for Traits 1 and 2, were sampled and plotted as points on the fitness surface, color-coded by generation, along with the mean population value plotted as the white line, to illustrate the adaptive walk of the population. (c-e) Aggregated across 500 simulations per set of parameters, as populations took adaptive walks, the effect size of fixed QTLs decreased geometrically. The mean allelic substitution effect size of each order of fixation is shown in black, overlaid on the mean founder population QTL. The geometric series *α_k_* = *α_1_a^k-1^* was fitted to each series of effect sizes, with the parameter estimates shown in each figure, and the dotted black line showing the line-of-best-fit for the fixation order. (f) A diagram of an international breeding network, in which breeding programs exchange germplasm. (g) The conceptual model of the evolution of crop germplasm and the origin of breeding metapopulations underlying our simulations. (i) The simulation pipeline began with the stochastic simulation of a founder population of 2000 individuals. (ii) Two ‘landrace’ populations were randomly selected from the founder population, and underwent independent recurrent selection for 200 generations. (iii) Landraces were independently selected and ‘purified’ through intensive selection followed by repeated selfing, generating two purelines per breeding program. (iv) Two different biparental purelines crosses were made: admixed (across breeding programs, assumed to be allo-elite) and unadmixed (within breeding program, be iso*-*elite). (v) A biparental RIL family from each cross with 250 lines was developed. (vi) The grouping of two breeding populations with germplasm exchange constituted a *breeding metapopulation*.

### Defining a breeding simulation as a history of a breeding program

#### Landrace phase

We first simulated a plausible history of the evolution of locally-adapted crop landraces. After 5-generations of burn-in of the founder population, two subpopulations (i.e. landraces) were created by randomly sampling without replacement (Fig. 1g(i)). Subpopulations 1 and 2 underwent 200 generations of independent recurrent selection, where at each generation, the top 10% of individuals were advanced based on *w*, and randomly mated to produce the next generation (Fig. 1g(ii)). This scenario was representative of two geographically isolated landraces being subject to the same environmental and cultural constraints. Since a complete absence of gene flow is expected to lead to greater genetic divergence, we partially counterbalanced this by not allowing for *de novo* mutation, to reduce the expected genetic divergence.

#### Breeding phase

Next, we simulated the origin of modern breeding metapopulations, in which local landraces were collected and purified by independent breeding programs, followed by intermating between different purified landraces. After the ‘landrace’ phase, each subpopulation was ‘purified’, passing through a bulk phenotypic selection breeding pipeline, which derived two purelines per subpopulation (Fig. 1g(iii)) (see Supplementary Table 5). Narrow-sense heritability was increased to 0.8 for the purification phase, and individuals were advanced on the basis of suitability alone. A biparental admixed recombinant inbred line (RIL) family was created by crossing one line from each independent subpopulation-derived pureline (Fig. 1g(iv) and 1g(v)). A biparental unadmixed RIL family derived from a cross between the two purelines derived from the same subpopulation also created as a negative control, by crossing the two purelines derived from the same subpopulation. QTL mapping was run on each of these RIL populations. For aggregated analyses, each breeding simulation was run 400 times with the same set of parameters.

### Linkage mapping

Single linkage mapping (Lander and Botstein 1989) was done with the scanone() function from the R package “qtl2” (Broman et al. 2019) using Haley-Knott regression (Haley and Knott 1992). Significance of LOD peaks was determined by running 1000 permutations to calculate a trait-specific, genome-wide Bonferroni correction at *α* = 0.05. The number of significant LOD peaks were counted based on LOD peaks exceeding these thresholds, with a support interval of 5. This conservative support interval, much higher than the standard 1.5 (Manichaikul et al. 2006), was selected to reflect the significant magnitude of the LOD peaks detected in this study from the high precision enabled by a high density of markers, and to allow for the counting of multiple LOD peaks per chromosome while minimizing the amount of false positive detections.

Epistatic (2-dimensional) linkage mapping (Wang et al. 1999) was done with the scantwo() function from the R package “qtl” (Arends et al. 2010) with Haley-Knott regression. Due to the computational runtime of running permutation tests for 2D QTL mapping, fixed thresholds of (6.6, 5.2, 4.2, 5.0, 3.1) for the full, conditional-interactive, interaction, additive, and conditional-additive LOD scores, respectively, were obtained, by running 10 independent 1000-fold permutation tests and taking the mean value for each threshold. For counting the number of significant interaction LOD peaks, only the highest interaction scores per chromosome were considered, for the sake of computational simplicity. This likely represented an underestimation of the total number of interactions, because additional peaks aside from the top peak on each chromosome were ignored.

### Principal component analysis

Principal component analysis (PCA) was done with the prcomp() function in base R (R Core Team 2025). The principal components were derived based on the genotypes of the two landrace populations. The admixed RIL family was then projected onto the first principal component, and color-coded by its parental haplotype with respect to the two QTL closest to the most significant interaction detected in epistatic mapping. The parental haplotypes were determined based on each of the two purelines derived from each of the two landraces that were used as parents in the biparental RIL cross.

### Quantification of genetic divergence

To quantify the genetic distance between purelines, an “isoeliteness” score was calculated as a normalized, weighted diploid Hamming distance of the two derived purelines with respect to the attained trait QTL:

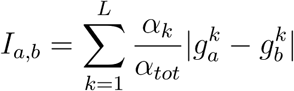

Where 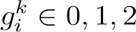 was the allelic dosage at QTL *k* for individual *i* (2 = homozygous for allele *0*, 0 = homozygous for allele *0* and 1 = heterozygous), and the weight term *α_k_ /* α*_tot_* was the allelic substitution effect at QTL *k* over the sum of allelic substitution effects across all QTL for that trait. The isoeliteness score fell between 0 (contrasting alleles at all QTL) and 1 (isogenic at all QTL). The “mean isoeliteness” was the average value across the two attained traits. The Hamming distance was calculated using the same equation, but without the weight term.

To quantify the degree of transgressive segregation, the “excess variance” for each trait was calculated as the change in phenotypic variance in the biparental RIL family relative to the phenotypic variance across both of the pureline populations from which a parent was sampled:

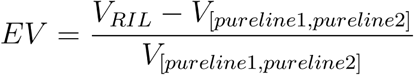

To quantify population divergence, an fixation index was calculated for the entire metapopulation as:

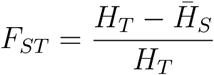

Where *H_T_* was the expected heterozygosity and *H̅_S_* was the mean expected heterozygosity across subpopulations, based on allelic frequencies and expectations under Hardy-Weinberg Equilibrium.

### Determining the aggregated geometric series of allelic effect sizes

Across 500 simulations, at each generation, the genotype of each QTL across each subpopulation were compared to that of the previous generation to determine whether it became fixed in that generation. A counter was incremented for each subpopulation at the end of each generation by the number of QTL that were fixed that generation, so that any QTL fixed in a given generation would have the same “order of fixation”. At the end of the simulations, the mean allelic substitution effect for each order of fixation was then calculated, discarding outliers at high orders of fixation with a sample size below 100. To fit a line-of-best-fit for the geometric series *α_k_* _=_ *α_1_a^k-1^* of the allelic fixation order, the nonlinear least squares (‘nls’) function with the “port” algorithm was used.

To determine the mean aggregated architecture for the founder populations and the admixed RIL families, the allelic substitution effects of each segregating QTL were sorted and ranked by decreasing effect size. The mean allelic substitution effect for each rank across the simulations was then calculated, and a slightly modified line-of-best-fit method compared to the order of fixed alleles was used, by adding an estimated intercept to account for the fact that the effect sizes in the founder trait architecture did not converge to zero. The estimate of *α_1_* for each admixed RIL family was projected onto the founder trait architecture by solving for *k* in the line-of-best-fit equation that was fit to the founder population from which the RIL was derived.

### Constructing a genotype-to-fitness landscape with the sorghum NAM

To investigate the fitness landscape dynamics of global breeding metapopulations using real data, we leveraged the sorghum nested association mapping (NAM) resource, as described in (Bouchet et al. 2017). This population consisted of 10 biparental F_6_ RIL families, each derived from crossing one of 10 diverse founder lines from the sorghum association panel (Casa et al. 2008) with a common parent (RTx430). For our analysis, we selected 5 families (SC265, SC283, SC35, Macia, and Ajabsido) based on the genotypic and phenotypic analysis in (Bouchet et al. 2017). Our selected families had a high degree of genetic divergence and varying levels of phenotypic similarity for the flowering time and plant height traits, the same traits the attained traits in our simulation were meant to represent. For each trait, we calculated the “optimal” value as the phenotype of the common parent, RTx430, as measured in Manhattan, KS in 2015. We applied the following normalization to give each trait value an optimum of zero and a reduced variance:

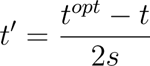

Where *t* was the original trait value, *t^opt^* was the optimal trait value, and *s* was the standard deviation for the trait across all families. This transformation enabled us to calculate suitability based on flowering time and plant height using the same suitability function as described above.

We next ran principal coordinate analysis of genome-wide polymorphism using genotype-by-sequencing (GBS) polymorphism data from the Sorghum Association Panel (SAP) (Morris et al. 2013), representing global sorghum diversity, projected the 5 selected NAM families onto principal components (PCs) 1 and 2, and color-coded each individual by suitability. Since there was a high level of noise among suitability values, we then smoothed the image by iteratively convolving a gaussian kernel with the smooth.2d() function from the R package “fields” (Nychka et al. 2025). We ran 15 smoothing iterations, clipping the resulting surface to be between 0 and 1 each time, and increasing the *theta* parameter from 1 to 15 to first filter out noise and then gradually aggregate spatial information to determine high-level features. We then used “plotly” to render the three-dimensional surface and overlay the NAM individuals. To recreate Wright’s original fitness landscape (Wright 1932), we ran a sliding scale with the base R focal() function to calculate local minima and maxima designated as the “peaks” and “valleys” of the landscape. Additionally, to determine the principal components that explained most of the variation for suitability, we fit a linear model with the lm() function to estimate both flowering time and plant height. We selected the principal component with the highest correlation for each trait to use for our alternative rendering of the fitness landscape (Supplementary Fig. 5).

## RESULTS

### Adaptive walks from standing genetic variation

To investigate the dynamics of adaptation on our fitness landscape, we tracked the genotypic and phenotypic responses of subpopulations as they took adaptive walks towards a fitness peak. By recurrently selected for breeding fitness (Fig. 1a), the desired trait (i.e. yield potential) was indirectly improved through directional selection, and the two attained traits (e.g. flowering time, plant height) were indirectly improved through stabilizing selection (Fig. 1b). These selection pressures were representative of crop landrace development, where sub-optimal suitability to an environment incurs a yield penalty, so selection for “breeding fitness” should lead to a gradual increase in adaptive suitability. In line with SBT, this adaptive walk was more direct in large populations than in small populations more impacted by genetic drift (Supplementary Fig. 1). By tracking the allelic frequencies over the course of each adaptive walk, we characterized the dynamics of adaptation from standing genetic variation. On average, the QTL with the largest effect size tended to get fixed first, followed by QTL with progressively smaller effect sizes, roughly following a geometric distribution (Figs. 1c-1e). This finding parallels the property of an exponential decrease in the size of fixed *de novo* mutations along an adaptive walk (Fisher 1930; Kimura 1983; Orr 1998) (Supplementary Table 4).

### Parallel adaptation of independent subpopulations

After two subpopulations were sampled from the same founder population and selected according to the same criteria, they exhibited phenotypic parallelism (Fig. 2a, Supplementary Fig. 2). The stochastic nature of their adaptive walks due to genetic drift and phenotypic selection was apparent in the allele frequencies of the QTL underlying the attained traits (Figs. 2b, 2c). The broad patterns of allelic frequency change in both subpopulations were similar, with large-effect QTL becoming fixed early on and smaller-effect QTL continuing to segregate, but the specific alleles fixed differed between subpopulations (Fig. 2d). Following independent adaptation, each subpopulation was purified through intensive selection for suitability, representative of the process by which nascent breeding programs developed their initial germplasm from locally-adapted landraces. From each subpopulation, two immortalized elite purelines were developed. On average, the isoeliteness score of purelines derived from different subpopulations, assumed to be allo-elite, was significantly lower (11% for *L* = 10 and 29% for *L* = 50) than the isoeliteness score of purelines derived from the same subpopulation, assumed to be iso-elite (Fig. 2e, Supplementary Table 6). Isoeliteness was not significantly correlated with genome-wide *F*_ST_ between the landrace subpopulations (Supplementary Fig. 3), indicating that genome-wide divergence is a poor indicator of genetic compatibility.

**Figure 2.**
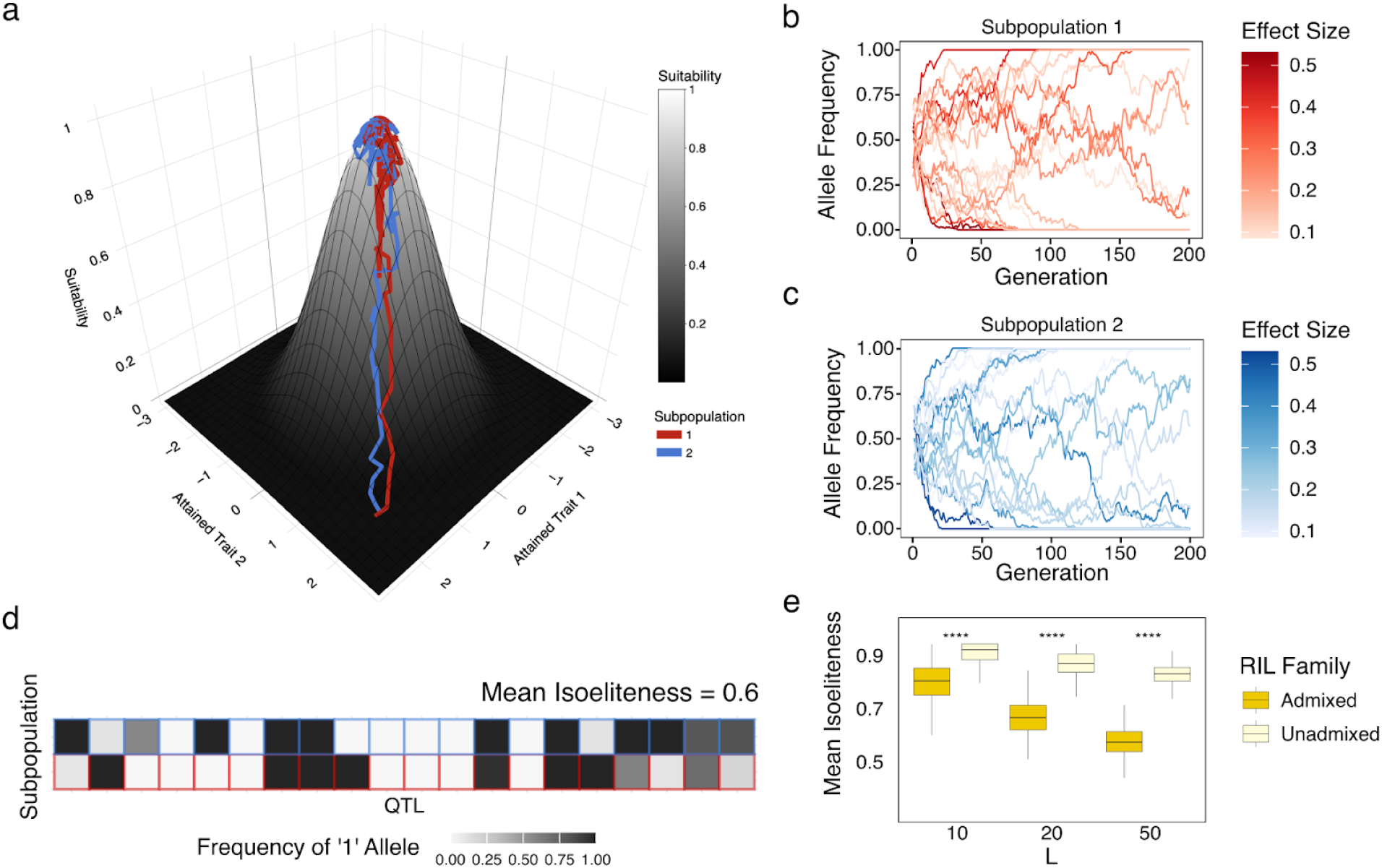
Stochasticity during local adaptation of independent subpopulations leads to cryptic genetic heterogeneity. (a) Two subpopulations were sampled from the same founder population (shown here is the *L* = 10 founder scenario), and took independent adaptive walks towards the fitness peak. The coordinates for the trajectory of each subpopulation were the mean attained trait values at each generation. (b and c) Allelic frequencies for both attained trait QTL of the ‘1’ allele were illustrative of the parallelism underlying genetic drift between the two subpopulations. QTL were shaded according to allele substitution effect size. (d) The allele frequencies for both attained traits in each subpopulation after 200 generations showed contrasting alleles are at or near fixation across the two populations at several loci, measured by mean isoeliteness across both attained traits. (e) Aggregated across 200 simulations per set of parameters, the mean isoeliteness between purelines derived from different subpopulations was significantly lower than the isoeliteness of purelines derived from the same subpopulation, in each trait architecture scenario. Isoeliteness decreased with a higher number of QTL per attained trait.

### Transgressive segregation in admixed RIL families

After landrace purification, purelines derived from different subpopulations were crossed to create biparental admixed RIL families, which were compared to unadmixed RIL families derived from two purelines derived from the same subpopulation. In the admixed families, crosses between parents allo-elite for attained traits produced excessive transgressive segregation (e.g. Fig. 3a), while crosses between parents that happened to be more iso-elite led to less transgression (e.g. Fig. 3b). On average, excess variance (*EV*) for the attained traits was higher in admixed families than in unadmixed families in all founder scenarios (Figs. 3f, 3g, Supplementary Table 6). Isoeliteness was highly correlated with *EV* (*r* = −0.89; Figs. 3k, 3l), but *F*_ST_ was not (Supplementary Fig. 3). There was a greater EV for suitability than for the attained traits (Figs. 3c, 3h), and suitability *EV* was highly correlated with mean isoeliteness (*r* = −0.89; Fig. 3m, Supplementary Fig. 3), but again *F*_ST_ was not (Supplementary Fig. 3). There was little *EV* for the desired trait (Fig. 3d), though *EV* was notably higher in admixed families (0.07–0.09) than in unadmixed families, where it was absent (Fig. 3i, Supplementary Table 6). Breeding fitness, a composite of suitability and yield potential, was highly transgressive (Figs. 3e, 3j, Supplementary Table 6), and its *EV* was also strongly correlated with mean isoeliteness (*r* = −0.88).

**Figure 3.**
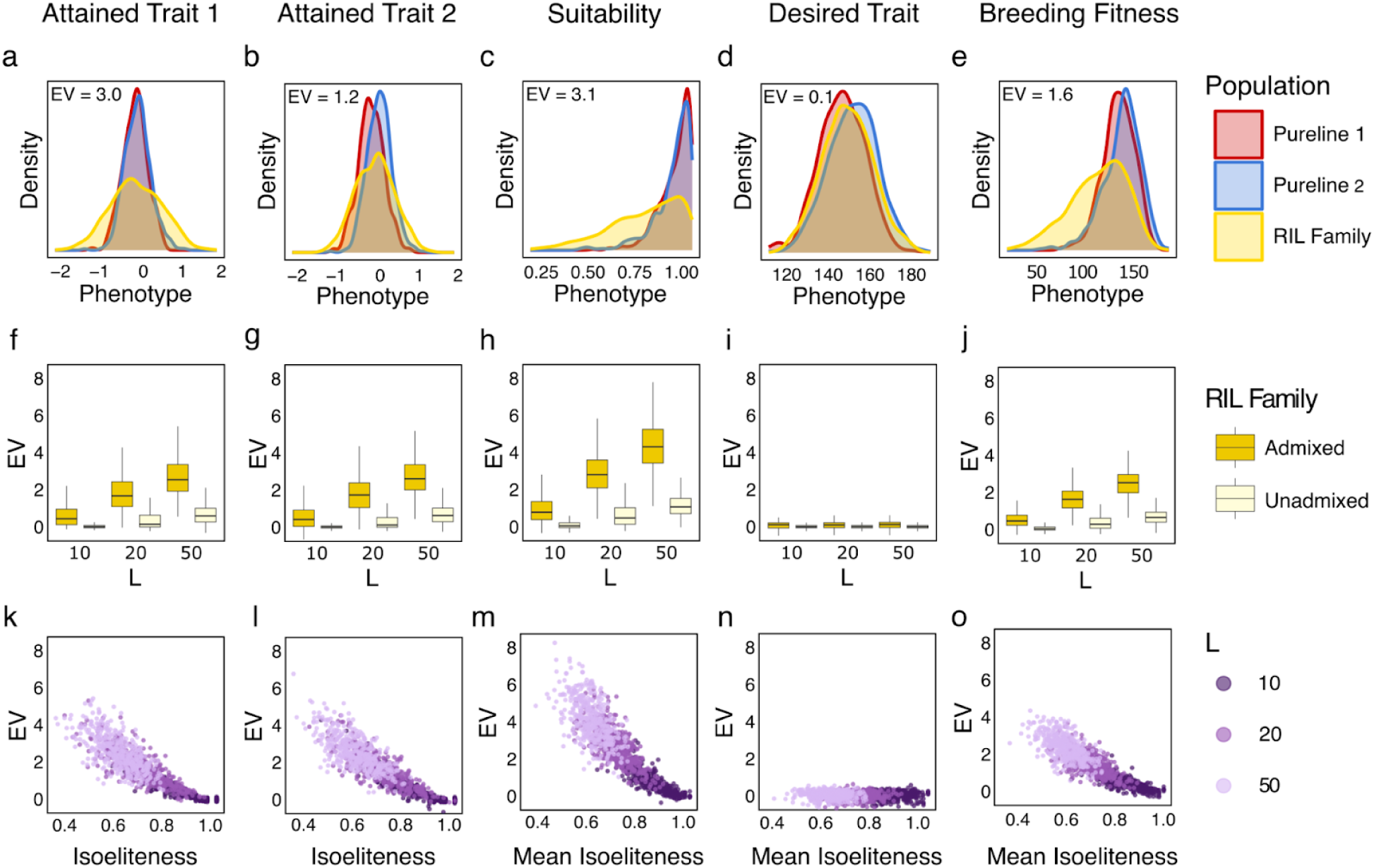
Admixture in allo-elite crosses leads to an excess of transgressive segregation. A cross between the purelines derived from the illustrative subpopulations in Figure 2a produced, there was excessive transgressive segregation for attained trait 1 ((excess variance) (*EV*) = 3.0) in the admixed RIL family (a), resulting from genetic heterogeneity between the pureline parents. There was less transgression for attained trait 2 (*EV* = 1.2), indicating more genetic homogeneity between the parents for large-effect QTL. Genetic heterogeneity across the two attained traits led to transgression for suitability (*EV* = 3.1) (c) and breeding fitness (*EV* = 1.6) (e). (d) The phenotypic distribution for the polygenic desired trait fell between the range of the two parents (*EV =* 0.1). Across 400 simulations per parameter set, there was significantly higher *EV* in admixed RIL families than in unadmixed RIL families for the attained traits (f, g), suitability (h), and breeding fitness (j). (i) There was a consistently low level of *EV* for the desired trait. Within the admixed families and aggregated across scenarios, isoeliteness was strongly correlated with *EV* for the attained traits (k, l), suitability (m), and breeding fitness (o). (n) Due to an absence of *EV* in the polygenic desired trait, and the independence with the attained traits, there was no correlation between mean isoeliteness and *EV* for the hidden desired trait.

### Architecture of attained traits and breeding fitness in admixed RIL families

We next ran linkage mapping on the RIL families to examine the genetic basis for the transgressive segregation. Linkage mapping of the attained traits detected more significant LOD peaks in admixed families than in unadmixed families across all founder scenarios (Figs. 4f and 4g, Supplementary Table 7). The number of peaks was highly correlated with isoeliteness (*r* =−0.85; Figs. 4k, 4l, Supplementary Fig. 3) and *EV* (*r* = 0.75; Supplementary Fig. 3). For the desired trait, there were a constant number of significant LOD peaks across all founder scenarios (Fig. 4i), though they were of much lower magnitude (Fig. 4d) than those of the attained traits (Figs. 4a, 4b), consistent with a polygenic trait architecture.

**Figure 4.**
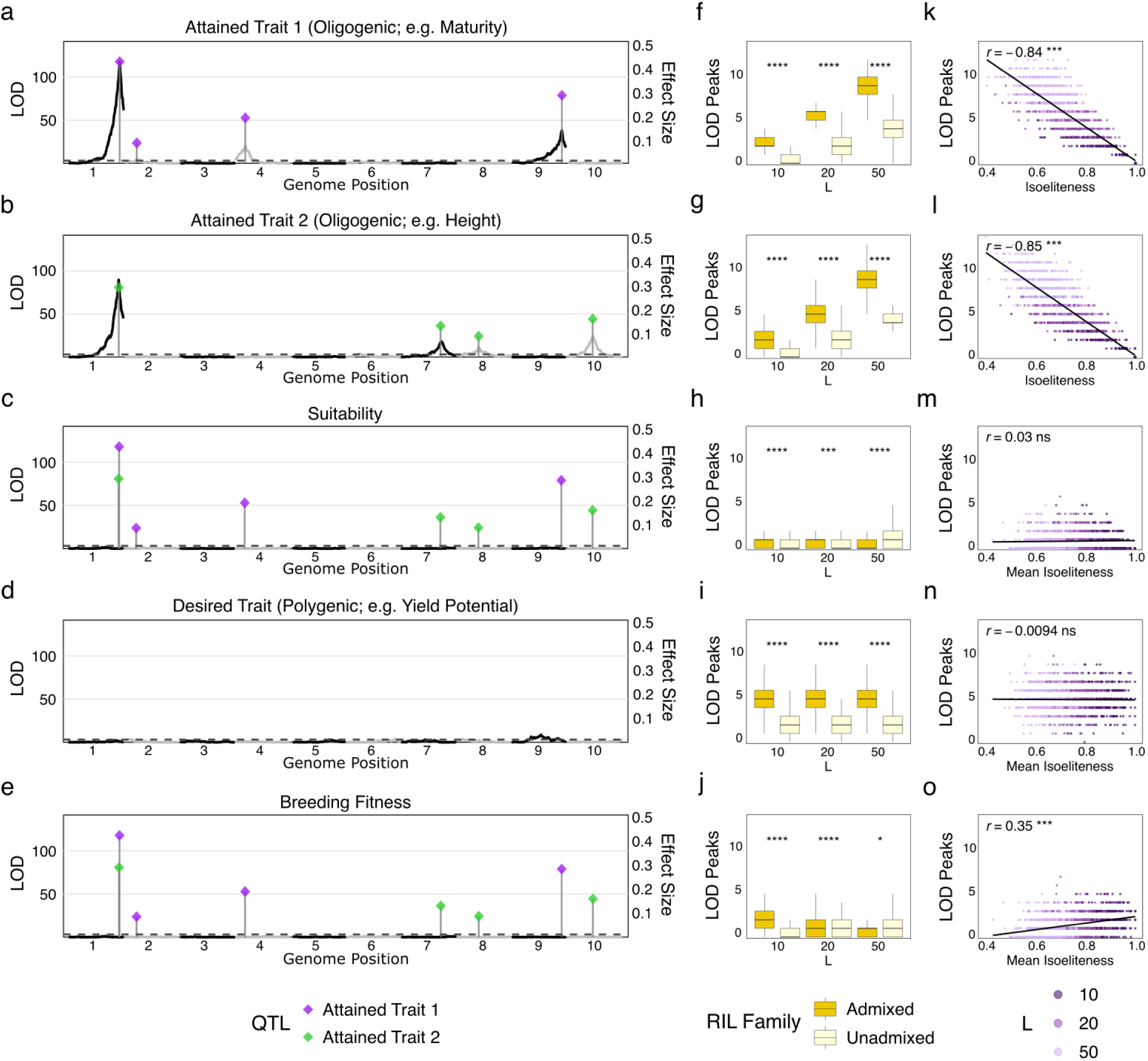
Release of genetic heterogeneity leads to an oligogenic trait architecture in admixed families. Linkage mapping was done on the admixed RIL family in the illustrative simulation, with significance thresholds determined by a genome-wide trait-specific Bonferroni correction at *α* = 0.05. Three significant LOD peaks (chromosomes 1, 3, 9) were detected for attained trait 1 (a) and four (chromosomes 1, 7, 8, 10) for attained trait 2 (b). These peaks were co-localized with the actual QTL (shown as diamonds), and correlated with allele substitution effect. No major peaks were detected for suitability (c) or breeding fitness (e). (d) There was one small desired trait LOD peak (chromosome 9), consistent with a polygenic trait architecture. (f, g) Aggregated across 400 simulations per set of parameters, there were more attained trait LOD peaks in the admixed RIL families than in the unadmixed RIL families, with a greater number of peaks detected in scenarios with higher *L*. More suitability peaks were detected in the admixed families in the *L* = 10 and *L* = 20 founder scenarios (h), but fewer in the *L* = 50 scenario. A consistent number of desired trait peaks were detected across all scenarios (i), and more breeding fitness peaks were detected in the admixed families in the *L* = 10 and *L* = 20 founder scenarios, but not in the *L* = 50 scenario (j). Within the admixed families, aggregated across scenarios, there was a strong correlation between isoeliteness and the number of attained trait peaks (k and l). There was a positive correlation between mean isoeliteness and breeding fitness peaks (o), and no correlation between mean isoeliteness and the number of significant peaks for suitability (m) or the desired trait (n).

For suitability, as with breeding fitness, there were notably few significant LOD peaks detected across all founder scenarios (Figs. 4h, 4j). Interestingly, there was a positive correlation (*r* = 0.35) between the number of breeding fitness LOD peaks and mean isoeliteness (Fig. 4o), indicating that segregation of attained trait QTL confounded identification of breeding fitness QTL. This was supported by the negative correlation (*r* = −0.32) between the number of attained trait and breeding fitness LOD peaks, the positive correlation (*r* = 0.18) between desired trait and breeding fitness LOD peaks (Supplementary Fig. 3), and the detection of fewer significant LOD peaks for breeding fitness in the *L* = 50 founder scenario than the *L* = 10 founder scenario (Fig. 4j).

On average, there was a geometric series of allelic substitution effects for the attained traits in the admixed families (Figs. 5a-5c), with a consistent mean *α_1_* (largest-effect QTL) across all founder scenarios (Supplementary Table 8). For each simulation, we projected the admixed RIL *α_1_* onto the scenario-specific initial trait architecture to determine the rank (*k*) of that QTL in the series of effect sizes in the founder population. This rank determined an approximate boundary between the number of QTL fixed during the “deterministic” and “stochastic” phases of selection, which decreased as the genetic complexity of the founder scenarios increased (*L* = 10: *k* = 5.0; *L* = 50: k = 2.5; Supplementary Fig. 4d, Supplementary Table 8). These thresholds approximated the number of large-effect QTL from the founder population that were fixed isogenically across the adaptive walks of both subpopulations before one was differentially fixed.

**Figure 5.**
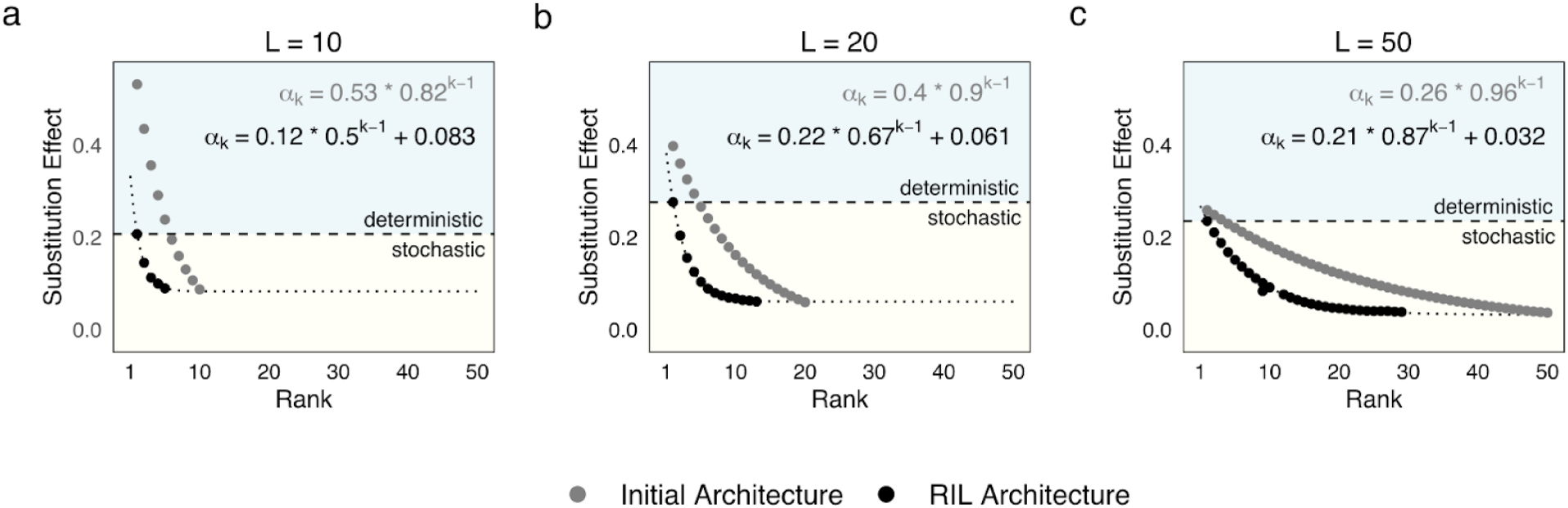
Selection produces a geometric series of effect sizes for attained traits in admixed families. The segregating QTL underlying variation in the attained traits in the admixed RIL families, contrasting the evolution of genetic architecture when starting from (a) 10 QTL, (b) 20 QTL, and (c) 50 QTL. The mean allelic substitution effect size of the ordered QTL in the admixed families, across 500 simulations per scenario, is shown in black, overlaid on the mean initial trait architecture (in grey) in each scenario. The geometric series *α_k_* _=_ *α_1_a^k-1^+ b* was fitted to each series of effect sizes, with the parameter estimates shown in each figure, and the dotted black line showing the line-of-best-fit for the aggregate RIL trait architecture. The intercept of *b* was added to account for the lowest possible effect size based on the founder series. The dashed line represents the approximate division between the QTL in the founder population that were typically fixed isogenically across the two subpopulations and those that were differentially fixed, and therefore segregated in the admixed RIL family.

### Fitness epistasis as an emergent property of stabilizing selection

We next ran epistatic linkage mapping on breeding fitness to test whether intergenic interactions were masking the effects of single QTL (e.g. Fig. 5e). On average, there were 1.6–4.2 times as many significant interaction LOD peaks in admixed families than in unadmixed families (Fig. 6d, Supplementary Table 7), and a strong correlation (*r* = −0.54) between the number of interactions and mean isoeliteness (Fig. 6e). Interestingly, the most significant interaction LOD peaks were found in the *L* = 20 scenario. These pairwise interactions (e.g. Fig. 6a) were due to crossover epistasis (e.g. Fig. 6b) between major QTL underlying the same attained trait (e.g. the QTL identified in Fig. 5a). This is because near a fitness peak, large-effect QTL will likely cause an individual to overshoot the fitness optimum, unless they are negated by an opposing QTL of similar magnitude.

**Figure 6.**
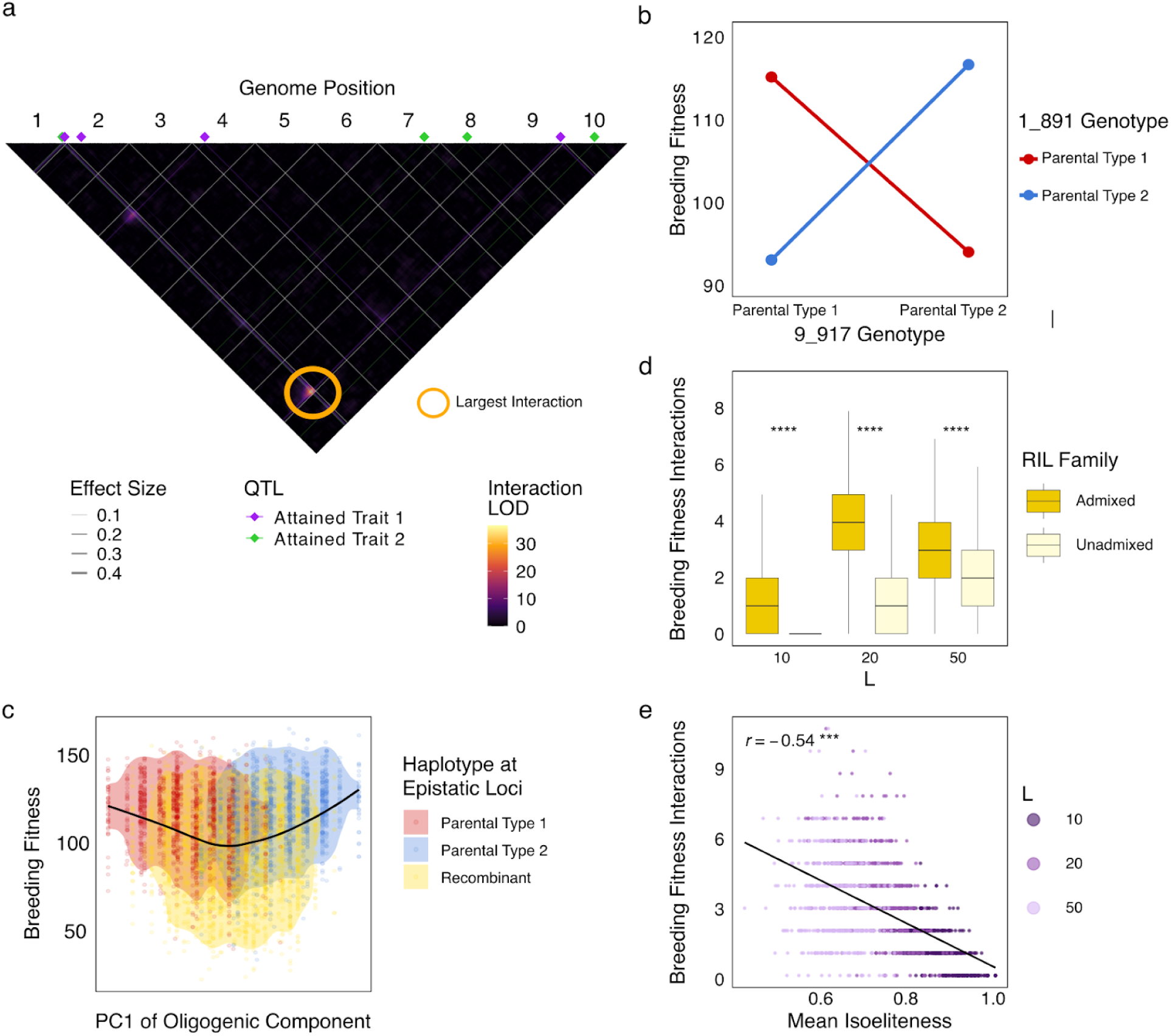
Epistatic linkage mapping reveals the emergence of fitness epistasis in admixed families. (a) Epistatic (2D) QTL mapping in the illustrative simulation run revealed a significant interaction effect between markers on chromosomes 1 and 9.The actual QTL, color-coded by trait, are shown as diamonds along the top and lines through the matrix, with widths of the bars determined by additive effect size. (b) A reaction norm plot with the two QTL nearest to the most significant interaction peak showed a crossover epistatic interaction, in which the effect on breeding fitness of one marker was dependent on the allelic state of the other marker. (c) PCA was run on the two subpopulations, and the admixed RIL family was projected onto PC1. Individuals were color-coded according to their parental haplotype, with respect to the same pair of markers from the most significant interaction peak. The mean breeding fitness along PC1 is shown in black. (d) Aggregated across 400 simulations per set of parameters, there were significantly more significant breeding fitness interactions in the admixed populations than in the unadmixed families across all scenarios. The most interactions were detected in the *L* = 20 scenario. (e) There was a strong correlation (*r* = −0.54) between mean isoeliteness and the number of significant breeding fitness interaction LOD peaks detected.

Running PCA on the unpurified landrace subpopulations and projecting the admixed RIL family onto PC1 produced a genotype-to-fitness landscape with two distinct peaks corresponding to each subpopulation (Fig. 6c). The individuals with the parental haplotypes of the QTL with significant interactions in the admixed family tended to be those with the highest fitness, while recombinant individuals had lower breeding fitness on average. Overall, fitness epistasis was a consistent emergent property of the attained trait genetic architecture in admixed families.

### An empirical genotype-to-fitness landscape in a global prebreeding population

We next applied our fitness landscape methodology using real data from the sorghum NAM resource, which consisted of 10 admixed RIL families, each derived from a cross between a diverse founder line and a common parent, RTx430, which was well-adapted to the U.S. environment where the resource was developed. We analyzed two attained traits: flowering time and plant height, and selected a set of phenotypically diverse families (Fig. 7a, 7b), of which three had narrow phenotypic distributions close to the optimum (SC35, SC283, Macia), and two had wide phenotypic distributions (Ajabsido, SC265). We then normalized the phenotypes and used our gaussian fitness function to estimate suitability, producing a range of suitability distributions from the most adapted family (SC35) to the least (Ajabsido) (Fig. 7c). A projection of the NAM onto PCs 1 and 2 of the SAP showed distinct clustering by family, with founders on the outer edge of each cluster (Fig. 7d).

**Figure 7.**
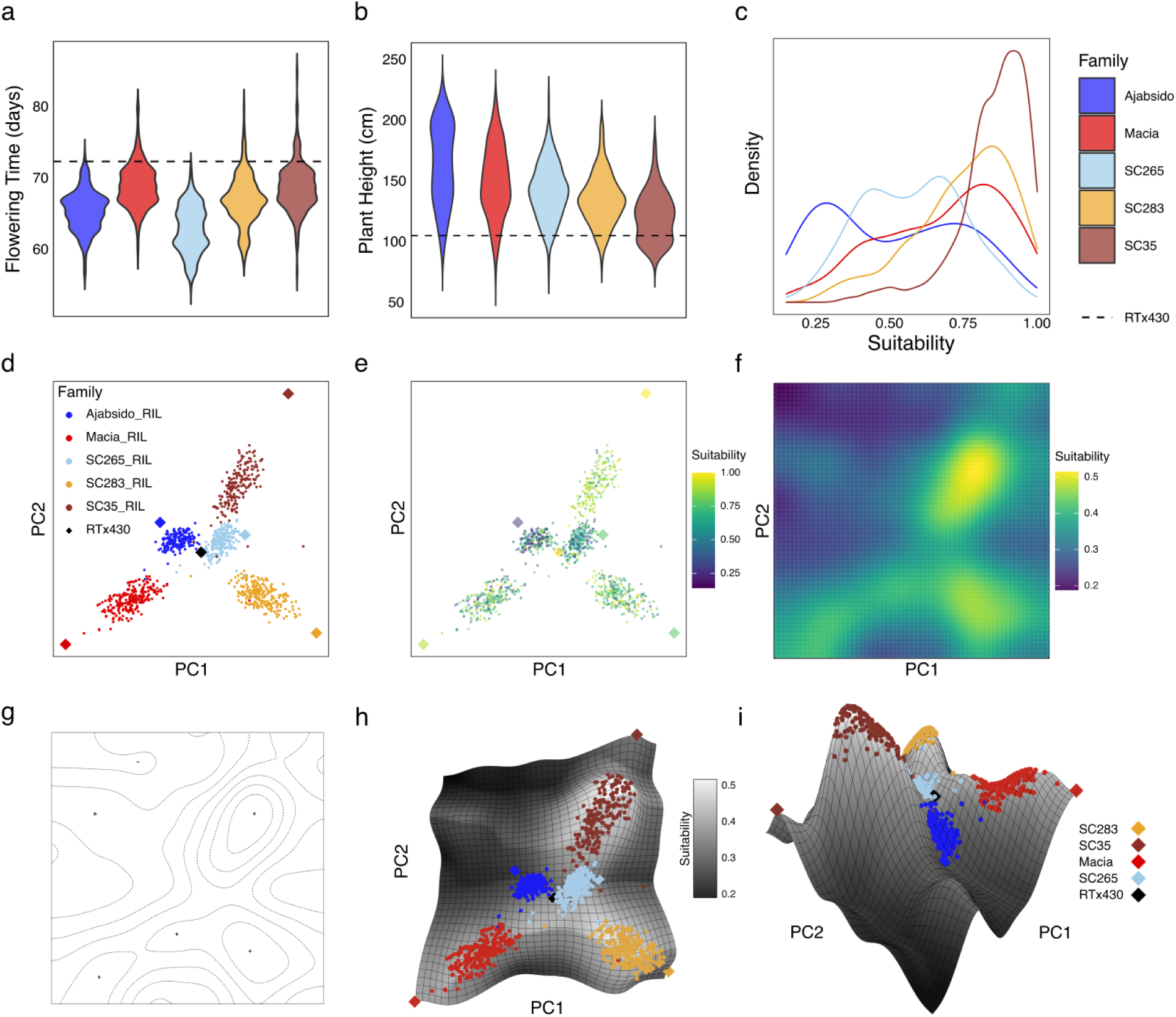
An epistatic genotype-to-fitness landscape exists in a real global prebreeding population. (a) The distribution of phenotypes for flowering time (a) and plant height (b) within each NAM family, along with the optimal phenotype (as determined by RTx430), showed that SC283, SC35, and Macia families were closest to the RTx430 phenotypes (shown as dashed lines), which was validated by the distribution of suitability values for each family (c). (d) Projecting the 5 families onto a PC1 and PC2 of the SAP showed a distinct clustering of families, with each diverse founder line on the edge of its family’s cluster, and RTx430, the common parent, near the center. (e) Each individual was color-coded by suitability, and iterative smoothing with a gaussian filter was applied to reduce noise and identify high-level spatial features of the fitness landscape to generate a smoothed landscape (f). (g) An empirical recreation of Wright’s canonical 1932 drawing of a fitness landscape, generated with the real sorghum NAM data. (h) Each individual, color-coded by family, was projected onto a 3D rendering of the fitness landscape. (i) A profile view of the landscape showed the SC35 family on the tallest fitness peak, with the SC283 family and Macia family on lower peaks. The SC35 parent and Macia parents were on the far left and right. The Ajabsido and SC265 families occupied separate fitness valleys separated by a saddle.

Color-coding individuals by suitability (Fig. 7e) showed noisy suitability patterns, which we refined with iterative gaussian smoothing (see Methods), (Fig. 7f). We then created an empirical fitness landscape in the style of Wright’s canonical hand-drawn diagram (Wright 1932) (Fig. 7g), and rendered a 3D surface from the empirical landscape, onto which we projected the NAM (Fig. 7h). Distinct topographic features were apparent, including three fitness peaks corresponding to the SC35, SC283, and Macia families, consistent with their relatively high suitability, and two fitness valleys occupied by the Ajabsido and SC265 families (Fig. 7i, Supplementary Fig. 4). The construction of this landscape, and its consistency with phenotypic analysis from (Bouchet et al. 2017) further validates the relevance of fitness landscapes in global breeding metapopulations.

## DISCUSSION

### Models of adaptive walks from standing variation are relevant for breeding programs

Fitness landscape studies typically use Fisher’s geometric model as a framework for the characterization of adaptation, which deals exclusively with the contributions of *de novo* mutations (Fisher 1930; Orr 1998; Gros et al. 2009; Martin 2014). This is rooted in the idea that QTL with large effects on fitness become rapidly fixed, and therefore will not remain in a population as standing variation (Kimura 1983). This assumption does not hold for crop improvement, in which fitness peaks are constantly shifting due to new environmental conditions or cultural preferences (e.g. Fig. 1b) (Allard 1999; Tester and Langridge 2010; Technow et al. 2026), and selectable genetic variation for rapid adaptation is limited to alleles already segregating or recently introgressed (Jain and Stephan 2017; Stetter et al. 2018; Wisser et al. 2019). This is particularly true of the nascent breeding populations that make up global public breeding networks (e.g. CGIAR-NARS networks; Fig. 1f), which were derived in recent decades from heterogeneous landraces (Dar et al. 2006; Walker and Alwang 2015; Fall et al. 2026).

In this study, we validated theoretical (Lande and Thompson 1990; Orr 1998; Yu et al. 2008) and empirical (Brown et al. 2011; Olatoye et al. 2020) expectations of a geometric distribution of effect sizes underlying complex traits. While a biological basis for this property remains unknown, the manner in which selection shaped trait architecture in metapopulations in our study suggests possible explanations. We can view the QTL fixed in the deterministic phase of selection (Fig. 5) as critical for a trait, and those fixed during the stochastic, or nearly neutral (Ohta 1992) phase, as more interchangeable. The simpler trait architecture in the *L* = 10 scenario may be representative of a highly conserved genetic pathway in which functional copies of a few core genes are required for an optimal phenotype, thereby constraining the genetic search space and leading to predictable purifying selection (Stern and Orgogozo 2009). The trait architecture in the *L* = 50 scenario instead may represent a genetic pathway with a high level of degeneracy (Barghi et al. 2020), due perhaps to large gene families or a flexible molecular pathway (Pickett and Meeks-Wagner 1995; Yeaman 2022).

The contrasting metapopulation genetic architectures from each scenario were also evident in the number of attained trait QTL identified through linkage mapping (Figs. 4f and 4g), consistent with the finding that increased genetic complexity reduces the likelihood of parallelism (Orr 2005). Future studies could characterize quantitative traits under stabilizing selection based on the complexity and degeneracy of their molecular pathways, and draw inferences on the expected genetic heterogeneity in diverse germplasm (Brown et al. 2011; Li et al. 2015; Bouchet et al. 2017; Wisser et al. 2019; Powell et al. 2022; Jiao et al. 2023; Zebell et al. 2025). Beyond the cereal examples used in this study, other examples of attained traits include those underlying yield components (Cassman 1994; Knowles and Knowles 2006; Park et al. 2014), and influencing fruit quality (Klee and Tieman 2018).

### Fitness epistasis is expected to be prevalent in breeding metapopulations

The generation of novel genetic diversity in breeding is often attributed to the disruption of “linkage blocks” from recombination, implying that fixed allelic combinations, either beneficial or detrimental, are physically linked (Hanson 1959; Miller and Rawlings 1967; Allard 1999; Jordan et al. 2005). We showed that favorable epistatic complexes need not be on the same chromosome, as segregation of QTL across the genome (Figs. 4a–b) in admixed families leads to breeding fitness epistasis (Fig. 6a). The fitness non-additivity of alleles (Fig. 6b) that were differentially fixed in each subpopulation (Fig. 2d) arose even though there was an additive genotype-to-trait map (Fig. 1a).

While the emergent property of fitness epistasis from stabilizing selection is well-understood (Wright 1935; Barton 1986; Whitlock et al. 1995; Martin et al. 2007; Gros et al. 2009) (Supplementary Table 3), our findings highlight the importance of this phenomenon specifically for breeding. For example, we showed how the limitations of single linkage mapping for detecting breeding fitness QTL (Fig. 4e) can be overcome with epistatic linkage mapping (Fig. 6a), and how a deleterious variant in an admixed population can be suppressed by another variant with which it co-evolved an epistatic framework (Fig. 6c). The purelines parents of the admixed RIL families were derived from populations that had distinct epistatic frameworks of moderate-to-large-effect QTL (Fig. 2d), which were recombined in the allo-elite crosses (Fig. 4e). To accelerate genetic gain, breeding programs must convert this epistatic variance to additive variance by re-constituting the favorable combinations of alleles (Allard 1999; Technow et al. 2021).

Through our characterization of genetic architecture in admixed families, we demonstrated that the SBT view of adaptation being driven by epistasis, genetic drift, and population subdivision (Wright 1932; Whitlock et al. 1995; Fragata et al. 2019) is relevant for breeding metapopulations. We validated empirical findings (Wisser et al. 2019; Wang et al. 2021) that when subpopulations are independently selected according to the same fitness constraint (Fig. 2a), the stochastic nature of drift (Figs. 2b, 2c) can lead them to the bases of different genetically-defined fitness peaks (i.e. Phase I of SBT), which they subsequently climb by fixing favorable allelic combinations (Fig. 2d) (i.e. Phase II of SBT). Later, when these subpopulations are recombined, their progeny may be transgressive for attained traits that were previously optimized (Figs. 3a, 3b), and fall into a fitness valley (Fig. 3e) (Fall et al. 2026).

Future studies could examine the robustness of our findings to different fitness functions (Gros et al. 2009) and genetic architecture parameters. The “genotype-to-fitness landscape” concept is often described as a useful metaphor (Stewart 2008; Messina et al. 2011; Technow et al. 2021), but our analyses of both simulated (Fig. 6c) and empirical (Fig. 7g) data show that it may also be considered in a literal sense, as previously proposed (Orr 1998). By constructing a one-dimensional fitness landscape from an admixed family, we demonstrated a mechanism by which recombinant progeny can cross from one fitness peak to another by retaining one of the favorable allelic combinations that underlies high fitness (Fig. 6c) (i.e. Phase III of SBT).

### Using fitness landscapes to optimize prebreeding and gene flow in global breeding networks

Transgressive segregation for breeding fitness (Fig. 3e) represents the risk of wasted resources associated with the exchange of breeding materials across different germplasms, since genotypes with suboptimal suitability are often discarded (Jordan et al. 2011; Fall et al. 2026). The inability to directly introduce exotic materials into elite genepools motivates the use of prebreeding populations to identify and introgress favorable alleles into elite genetic backgrounds (Hallauer and Sears 1972; Holley and Goodman 1988). Prebreeding unadapted germplasm can be likened to intentional application of Phase III of SBT, in which favorable gene combinations from highly-adapted subpopulations spread to other subpopulations (i.e. “conversion”) and allow subpopulations with lower fitness to traverse fitness valleys. Thus, a small number of favorable interacting alleles may form a “bridge” in the fitness landscape.

An example of this “bridge” is the Sorghum Conversion Program (SCP) from the 20th century, in which early maturity (*ma1*) and semi-dwarfing (*dw1*, *dw2*, *dw3*) loci were introgressed into exotic landraces, facilitating effective geneflow from those landraces into U.S. breeding populations (Stephens et al. 1967; Jordan et al. 2011; Morris et al. 2013; Thurber et al. 2013). Like most prebreeding initiatives in the 20th century, the SCP was carried out with phenotypic selection, so it was slow, labor-intensive, and not an optimal means of constructing optimal haplotypes (Stephens et al. 1967; Thurber et al. 2013). Through genomic mapping and improved phenotyping tools, we can now identify oligogenic QTL underlying attained traits (e.g. Figs. 4a-b) and putative epistatic interactions (e.g. Fig. 6a) to improve marker-assisted selection and accelerate prebreeding efforts (Lande and Thompson 1990; Bouchez et al. 2002; Cobb, Biswas, et al. 2019). Given the predictiveness of isoeliteness for excessive transgressive segregation (Fig. 3o) and fitness epistasis (Fig. 6e), this metric would be valuable for designing crosses between parents across breeding populations and identifying parental haplotypes in the progeny of allo-elite crosses.

The presence of distinct peaks and valleys along the PCs underlying polygenic variation on the empirical NAM fitness landscape is the “ruggedness” that is expected to emerge on a genotype-to-fitness landscape with a high degree of epistasis (Kauffman and Levin 1987; Cooper and Podlich 2002; Hwang et al. 2017; Technow et al. 2021), in contrast to the smooth trait-to-fitness landscape we rendered for our simulations (Fig. 1b). When we instead projected the NAM onto the PCs underlying the most variation for flowering time and plant height specifically, the resulting fitness landscape was a less-rugged surface with the well-adapted families clustered together on one peak (Supplementary Fig. 4m). This surface was a clearer reflection of the favorable epistatic complexes underlying high fitness than the surface defined by the PCs of polygenic variation. Furthermore, the existence of a single flowering time peak (Supplementary Fig. 4j) and two plant height peaks (Supplementary Fig. 4k) may provide insight into the degree of genetic heterogeneity underlying each of those traits. This ability to leverage the hyperdimensionality of the genotypic space to generate hypotheses about genetic compatibility within metapopulations underscores the value of an empirical fitness landscape for global crop improvement efforts.

## DATA AVAILABILITY

Code and data is published to Github at https://github.com/CropAdaptationLab/FitnessLandscapes.

## ACKNOWLEDGEMENTS

GPM’s contribution to this study was supported by NSF Award #2421208 “Collaborative Research: EDGE CMT: New methods to elucidate the omnigenic model and other emerging concepts in complex trait genetics”. Support was also provided by the Gates Foundation through the grant “Green Evolution—Accelerating Dryland Cereals Improvement for Africa (INV-053669)” to GPM. TM’s graduate research assistantship was funded by the project “Artemis: Imaging Technology for a Food Secure Future (INV-068946)” led by Alliance of Bioversity & CIAT and supported by the Gates Foundation.

## SUPPLEMENTARY INFORMATION

**Supplementary Table 1.** References to fitness landscapes as a metaphor.

| Authors | Year | Definition |
| --- | --- | --- |
| Wright | 1935 | "It has seemed to me (1929 et seq.) that the central problem of evolution (as of live stock improvement (Wright, 1920, 1922)) is that of a trial and error mechanism by which the locus of a population may be carried across a saddle from one peak to another and perhaps higher one. This view contrasts with the conception of steady progress under natural selection developed in most extreme form by Fisher (1930)." (Wright 1935) |
| Lande | 1976 | "The value of an adaptive topography is that it is easily visualized and so makes the evolutionary dynamics of the population intuitively clear" (Lande 1976) |
| Orr | 1998 | "Indeed it may even be possible to decompose any real organism, which comprises an unspecifiable number of arbitrary characters, into an ideal one having some "effective" number of orthogonal characters, although I do not pursue this here." (Orr 1998) |
| Podlich DW,<br>Cooper M | 1999 | "Therefore, with the Wright model it can be argued that breeding programs should operate to exploit local peaks by selecting specific desirable epistatic combinations of genes but at the same time maintain a capacity to search for new higher peaks. Fisher's model, considered in terms of Wright's landscape concept, is a simplification of the shape of the landscape which assumes a single peak." (Podlich and Cooper 1999) |
| Allard | 1999 | "[Composite Cross II] remained highly variable genetically, and its multilocus organization appeared to fluctuate about a rarely, if ever, realized "adaptive peak" corresponding to the genetic structure appropriate to the weighted means of the environmental conditions that had prevailed during many preceding generations" (Allard 1999) (pp. 146) |
| Stewart | 2008 | "Although this specific theory [SBT] is still debated, it is based on two concepts that are highly relevant to plant breeding: fitness surfaces and adaptive peaks" (Stewart 2008) (pp. 54-55) |
| Messina | 2011 | "The implications of rugged landscapes for plant breeding are consequential as they suggest trajectories of genetic improvement conditioned by the presence of hills (high performance) and valleys (low performance)." (Messina et al. 2011) |
| Technow | 2021 | "As a metaphor it [the genetic landscape] should not be taken literally but can help gain an intuition for the complexity and ruggedness..." (Technow et al. 2021) |

**Supplementary Table 2.**
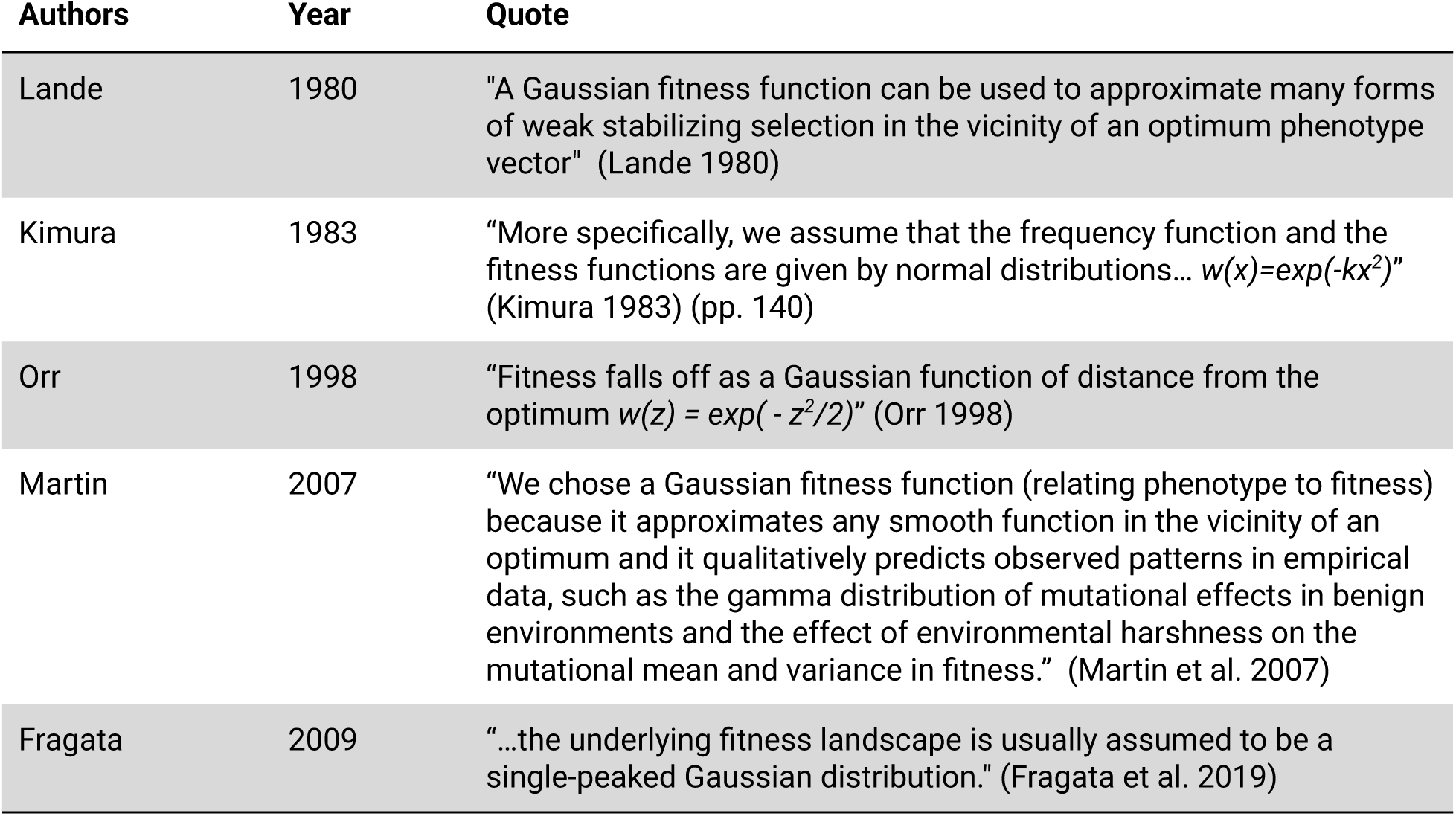
References to literature definitions of a phenotype-to-fitness landscape.

**Supplementary Table 3.** References to emergent fitness epistasis.

| Authors | Year | Quote |
| --- | --- | --- |
| Wright | 1935 | "The system of peaks relative to one character is not independent of that relative to another. Moreover, it is the harmonious adjustment of all characteristics of the organism as a whole that is the object of selection, not the separate metrical 'characters' " (Wright 1935) |
| Kauffman and Levin | 1987 | "In more realistic circumstances of strong epistasis in a complex genomic system, with intermediate levels of linkage and recombination, the consequent genetical constraints, mirrored in the vigorous lability of phenotypic properties as the genome changes, guarantees that the fitness landscape will be very rugged, with many peaks, ridges and valleys." (Kauffman and Levin 1987) |
| Whitlock | 1995 | "Stabilizing selection on polygenic traits, even when the genetic effects and variance for the phenotype are additive or involve dominance, generates epistasis for fitness" (Whitlock et al. 1995) |
| Brodie | 2000 | "Thus, the fitness effects of loci can be context dependent even when the phenotypic effects are strictly additive...*Whenever selection is nonlinear, loci with additive phenotypic effects will have nonadditive fitness effects.*" (Wolf et al. 2000) |
| Martin | 2007 | "Any model of stabilizing selection (selection for a given optimum) naturally generates epistasis for fitness, even when mutations act additively on the underlying phenotype, because the relationship between phenotype and fitness is nonlinear, as in our model here" (Martin et al. 2007) |
| Kopp and Hermisson | 2009 | "These alleles interact with each other, because stabilizing selection entails epistasis for fitness: Each mutant allele that increases in frequency brings the population mean closer to the optimum and, thereby, decreases the selection coefficient of the other alleles." (Kopp and Hermisson 2009) |
| Gros | 2009 | "It has been suggested for a long time that epistasis could emerge from the phenotype to fitness relation in an integrated model of phenotypic adaptation with continuous effects, referred to as the Fisher geometric model." (Gros et al. 2009) |
| Fragata | 2019 | "Mutations in FGM are usually assumed to be additive at the phenotype level. Nevertheless, mutational effects become epistatic at the fitness level because the relationship between phenotype and fitness is non-linear; the underlying fitness landscape is usually assumed to be a single-peaked Gaussian distribution." (Fragata et al. 2019) |
| Bank | 2022 | "Interestingly, the single-peaked phenotype-fitness landscape of FGM can give rise to an underlying genotype–fitness landscape with multiple peaks" (Bank 2022) |

**Supplementary Table 4.** What is known vs novel for geometric series in adaptive walks.

| Authors | Year | Finding |
| --- | --- | --- |
| Kimura | 1983 | The first allele fixed along an adaptive walk is likely to be a mutation of moderate effect size. (Kimura 1983) |
| Orr | 1998 | There is an exponential distribution in the size of mutations that become fixed along an adaptive walk. (Orr 1998) |
| Orr | 2005 | The frequency of parallel evolution declines based on the number of segregating mutations. (Orr 2005) |
| Kopp and Hermisson | 2009 | Standing variants of moderate effect are likely to be fixed first in adaptation to a moving optimum. (Kopp and Hermisson 2009) |
| This study |  | The order of standing variants fixed along an entire adaptive walk decreases geometrically. |
| This study |  | The segregating standing variation underlying a complex trait follows a geometric series of effect sizes – suggested by (Brown et al. 2011). |

**Supplementary Table 5.** Landrace purification via bulk selection approach.

| Population | Population Size | Action |
| --- | --- | --- |
| F1 | 1000 | Landrace population was randomly mated |
| F2 | 4000 | F1 population was selfed and 4 progeny per individual were advanced |
| F3 | 4000 | Top 2000 F2s are selected, selfed, and 2 progeny per individual were advanced |
| F4 | 2000 | Top 1000 F3s are selected, selfed, and 2 progeny per individual were advanced |
| F5 | 1000 | Top 500 F4s are selected, selfed, and 4 progeny per individual were advanced.<br>There were 2 F5 populations derived from the F4 population, to generate 2 different purelines. Each F5 had 2 progeny from each F4 line. |
| F6 | 500 | Top 250 F5s were selected, selfed, and 2 progeny per individual were advanced |
| F7 | 250 | Top 125 F6s were selected, selfed, and 2 progeny per individual are advanced |
| F8 | 240 | Top 60 F7s were selected, selfed, and 4 progeny per individual are advanced |
| F9 | 120 | Top 30 F8s were selected, selfed, and 4 progeny per individual are advanced |
| F10 | 60 | Top 15 F9s were selected, selfed, and 4 progeny per individual are advanced |
| Pureline | 1 | The top F10 individual was selected |
| Pureline Population | 500 | The pureline seed was bulked and selfed for 8 generations to immortalize it |
| Biparental RIL Family | 1000 | Purelines were crossed to create 250 full-sibling F1s. Each F1 was selfed to create 4 progeny, and each of those lines was selfed for 9 generations to create RILs. |

**Supplementary Table 6.** Breeding simulation aggregated results.

| QTL | Cross Type | Isoeliteness (Attained) | Isoeliteness (Desired) | Variance Expansion (Attained Traits) | Variance Expansion (Suitability) | Variance Expansion (Desired Trait) | Variance Expansion (Breeding Fitness) |
| --- | --- | --- | --- | --- | --- | --- | --- |
| 10 | Admixed | 0.86 | 0.82 | 0.59 | 0.90 | 0.07 | 0.53 |
| 10 | Unadmixed | 0.96 | 0.94 | 0.12 | 0.19 | 0.01 | 0.12 |
| 20 | Admixed | 0.72 | 0.82 | 1.71 | 2.75 | 0.07 | 1.58 |
| 20 | Unadmixed | 0.93 | 0.94 | 0.37 | 0.61 | 0.01 | 0.37 |
| 50 | Admixed | 0.63 | 0.82 | 2.58 | 4.16 | 0.09 | 2.41 |
| 50 | Unadmixed | 0.89 | 0.94 | 0.68 | 1.12 | 0.00 | 0.67 |

**Supplementary Table 7.**
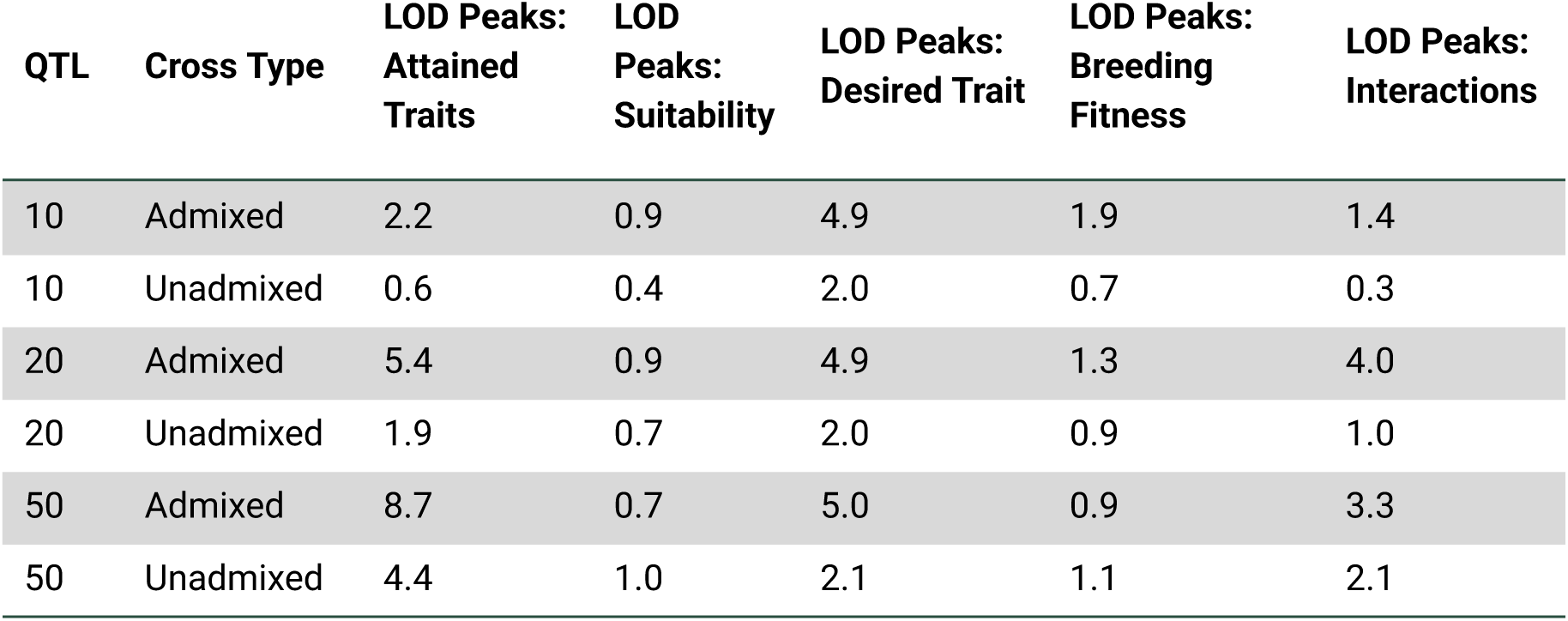
Linkage mapping aggregated results.

**Supplementary Table 8.** Geometric series of allelic substitution effects aggregated results.

| QTL | Founder alpha_1 | Founder alpha variance | RIL alpha_1 | RIL alpha variance | Mean Rank first Allogenic QTL |
| --- | --- | --- | --- | --- | --- |
| 10 | 0.53 | 2.2e-02 | 0.19 | 5.3e-03 | 5.0 |
| 20 | 0.40 | 1.1e-02 | 0.28 | 7.1e-03 | 3.8 |
| 50 | 0.26 | 4.2e-03 | 0.24 | 3.7e-03 | 2.5 |

**Supplementary Figure 1.**
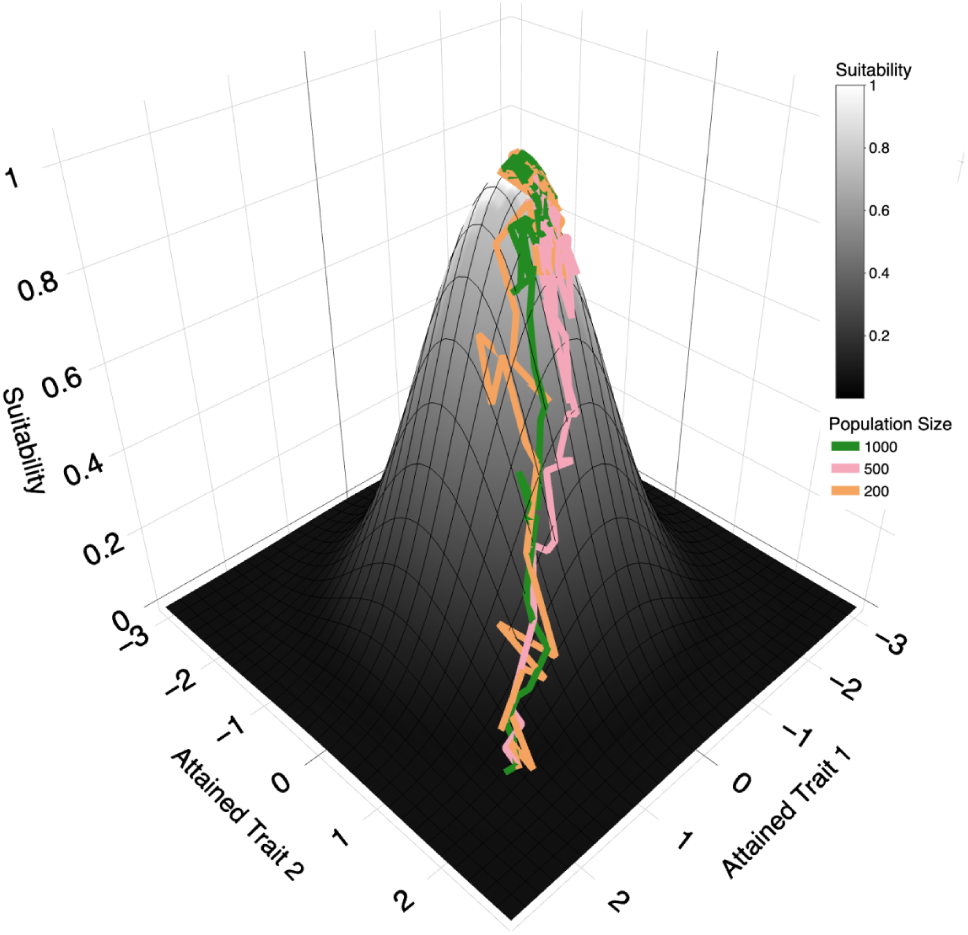
Adaptive walks of populations of different sizes. Populations of 1000, 500, and 200 individuals were selected for breeding fitness, taking adaptive walks towards the fitness optimum. Selection was the most effective on the large population, evidenced by its direct path to the peak. Selection was less effective on the small population, as it was more affected by genetic drift.

**Supplementary Figure 2.**
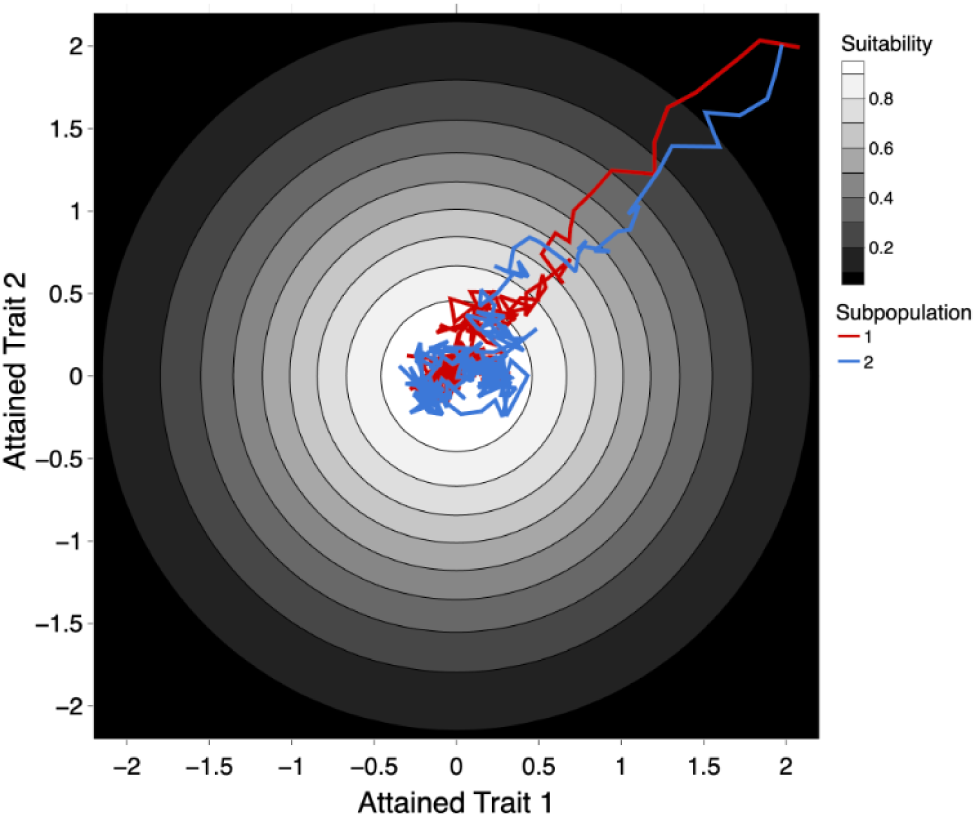
Parallel adaptive walks on a 2D contour map. An alternative, two-dimensional perspective of the same parallel adaptive walks depicted in Fig. 2a.

**Supplementary Figure 3.**
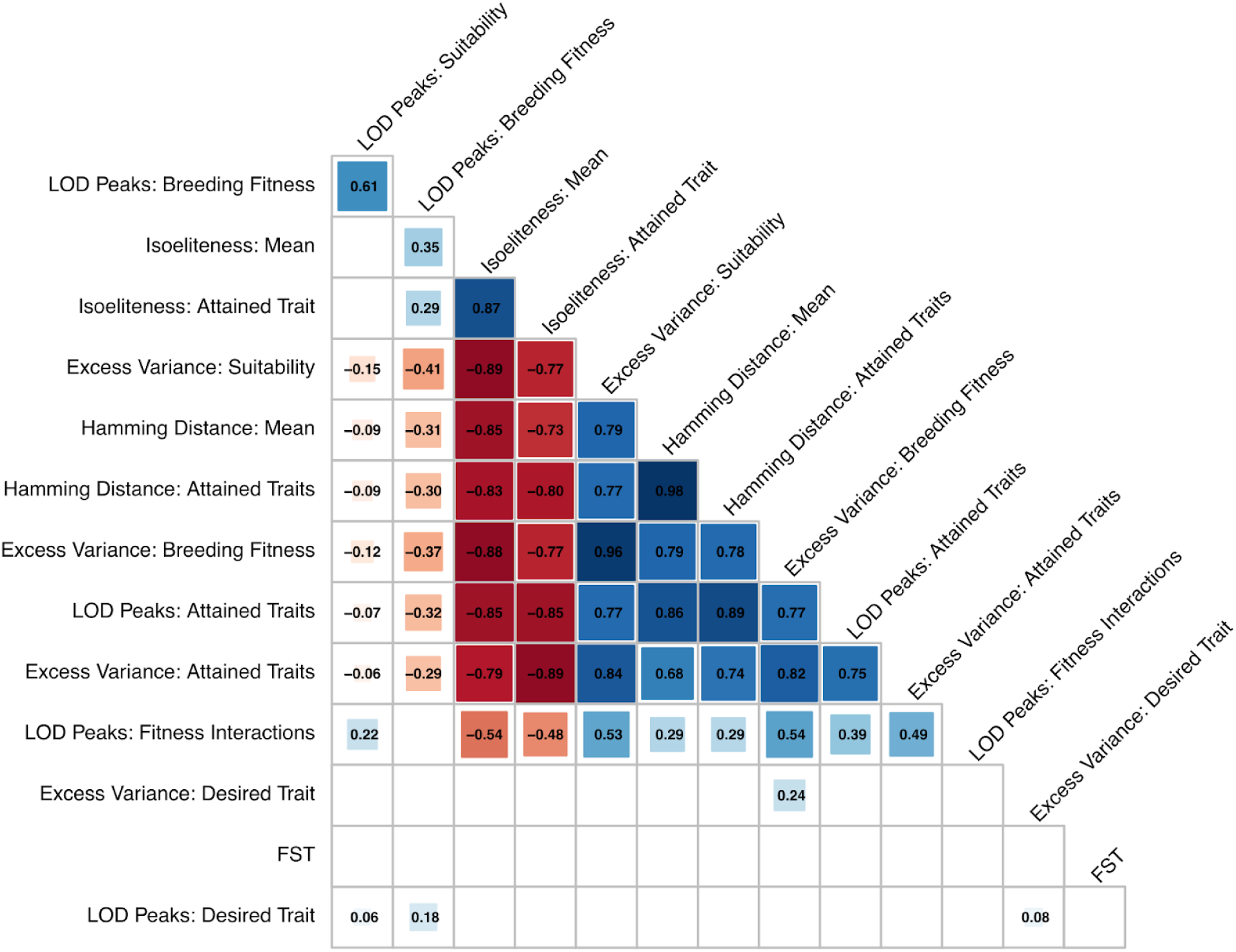
Correlation matrix between measures of genetic similarity and linkage mapping results. Shown here are the correlations, aggregated across all scenarios, of the metrics used to quantify genetic similarity (isoeliteness), phenotypic similarity (excess variance), and statistical relationships between markers and phenotypes (LOD peaks).

**Supplementary Figure 4.**
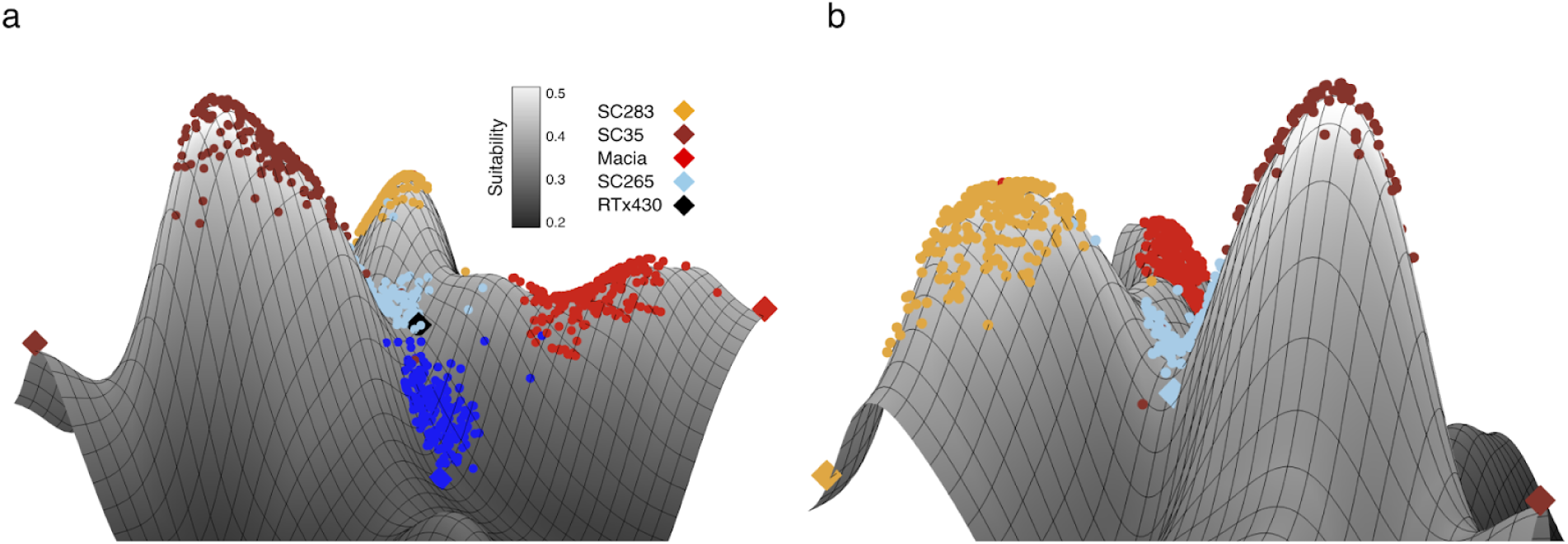
Perspectives of the sorghum NAM on a genotype-to-fitness landscape. As described in Figure 7, an empirical genotype-to-fitness landscape was created with the sorghum NAM resource. (a, b) Three fitness peaks and two fitness valleys were present on the landscape. The high fitness in the SC35 and SC283 families, both derived from parents from the Sorghum Conversion Program, and the Macia family, which was derived from the same zera-zera caudatum gene pool as RTx430 (Bouchet et al. 2017), was reflective of their adaptedness to U.S. environments, and the putative isoeliteness of each of the parents for flowering time and plant height with the elite US breeding line RTx430. Each of the founder lines of the families with high suitability sat on the edge of the landscape, in a region of lower fitness, demonstrating the value of a prebreeding program for improving the suitability of breeding lines (Jordan et al. 2011). Unlike the high-performing families, Ajabsido, an unconverted line, and SC265, a low-performing sorghum conversion line, resided in fitness valleys. SC265 may not have been fixed for a favorable combination of flowering or plant height alleles, but this would require further investigation. The low suitability of RTx430, which would be expected to have optimal fitness, was due to its high polygenic similarity with SC265 (Figure 7c). The low fitness in the SC265 family depressed the fitness landscape in the genetic space around RTx430, reflecting the fact that our fitness landscape was more useful for examining broad spatial patterns than the fitness of individuals.

**Supplementary Figure 5.**
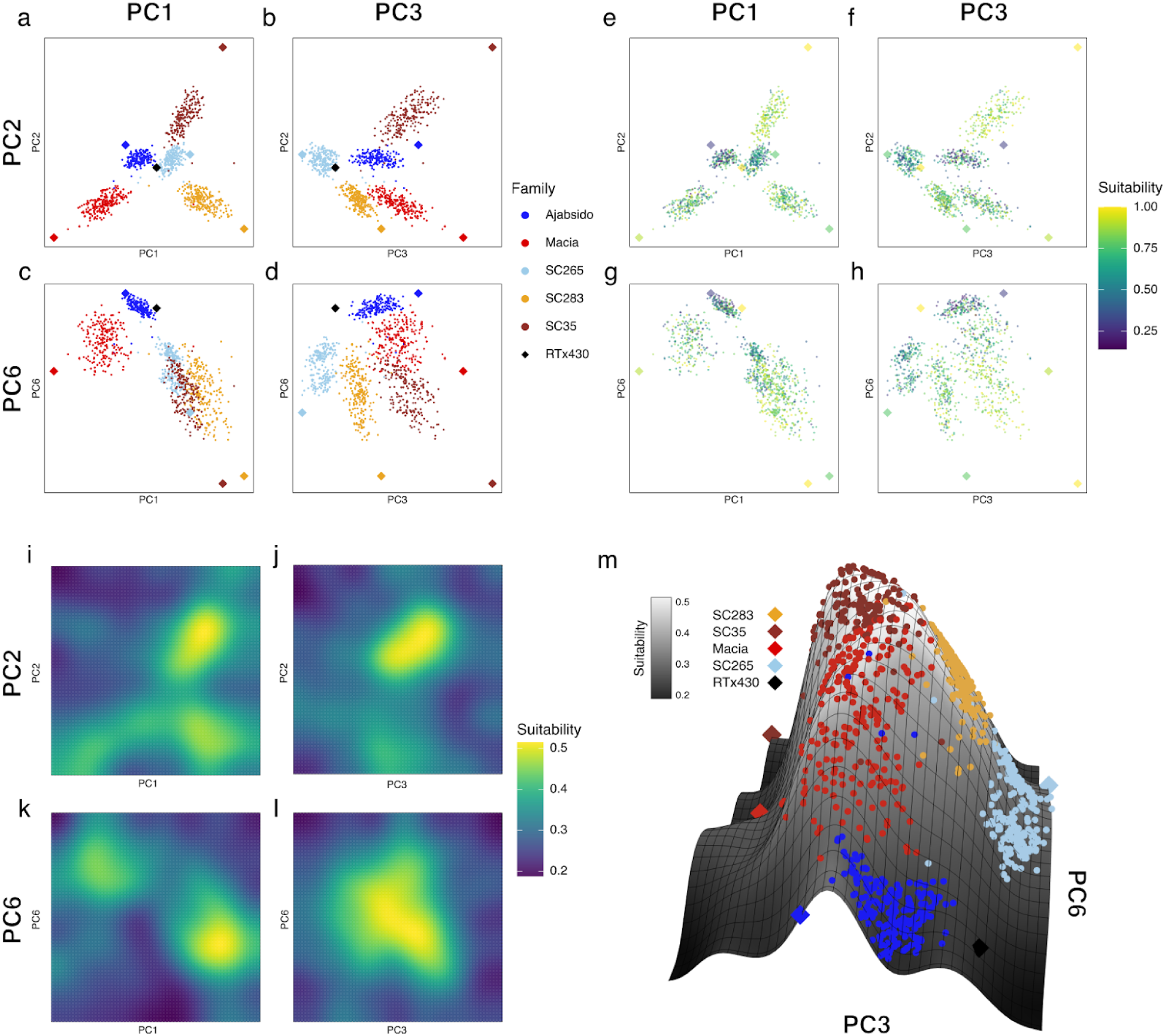
Projecting the sorghum NAM on the PCs underlying the most variation in suitability. Linear models were fitted to the flowering time and plant height phenotypes to determine the principal component that explained the highest amount of variation for each trait. PC3 explained the most flowering time variation and PC6 explained the most plant height variation. “Stretching” PC1 and PC2 (a) along these axes revealed clusterings of families by the genetic factors underlying flowering time (b) plant height (c), and both traits (d). This clustering demonstrated more apparent spatial patterning for suitability (f, g, h) among the families than was apparent along PCs 1 and 2 (e). Smoothing of the surface revealed a single peak for flowering time (j), two peaks for plant height (k), and a single peak for suitability (l). (m) A 3D surface rendering of the surface defined by PC3 and PC6 showed all 5 populations on the same fitness peak.

